# Adaptive Tree Proposals for Bayesian Phylogenetic Inference

**DOI:** 10.1101/783597

**Authors:** X. Meyer

## Abstract

Bayesian inference of phylogenies with MCMC is without a doubt a staple in the study of evolution. Yet, this method still suffers from a practical challenge identified more than two decades ago: designing tree topology proposals that efficiently sample the tree space. In this article, I introduce the concept of tree topology proposals that adapt to the posterior distribution as it is estimated. I use this concept to elaborate two adaptive variants of existing proposals and an adaptive proposal based on a novel design philosophy in which the structure of the proposal is informed by the posterior distribution of trees. I investigate the performance of these proposals by first presenting a metric that captures the performance of each proposals within a mixture. Using this metric, I then compare the adaptive proposals performance to the performance of standard and parsimony-guided proposals on 11 empirical datasets. Using adaptive proposals led to consistent performance gains and resulted in up to 18-fold increases in mixing efficiency and 6-fold increases in converge rate without increasing the computational cost of these analyses. [Bayesian inference; Adaptive tree proposals; Markov chain Monte Carlo; phylogenetics; posterior probability distribution.]

## Introduction

Studies relying on Bayesian inference of phylogenies are routinely conducted thanks to the numerous software packages designed for this purpose (Yang and Rannala 1997, 2012). These packages implement the Markov chain Monte Carlo algorithm (MCMC) to estimate the posterior distribution of parameters of a model capturing the evolutionary history (phylogeny) of taxa and their mode of evolution. Despite the pervasiveness of these analyses, estimating such posterior distributions remains a computational challenge whose complexity largely results from difficulties exploring and sampling the space of tree topologies.

The challenge of exploring the tree space has been recognized since the earliest days of Bayesian phylogenetic inference (Huelsenbeck et al. 2001). Long analyses and failure to explore the region of high posterior probability was shown to be a common occurrence that increased in frequency with the number of taxa studied (Beiko et al. 2006). Large datasets (taxa-wise) frequently resulted in rugged posterior distributions where clusters of tree topologies with high posterior probabilities were separated by low-probability valleys. A decade ago, the use of Metropolis-coupled MCMC algorithm (MC^3^; Altekar et al. 2004) was proposed as a solution to this major issue and, although this solution is now considered as a standard practice, the settings required for this method to perform appropriately remains a practical concern (Whidden and Matsen 2015; Brown and Thomson 2018). Using the MC^3^ algorithm reduces the failure rate by easing the sampling of rugged tree posterior distribution, but is not significantly improving the sampling efficiency of the key actors of the exploration of the tree space: the tree proposals.

In the last decades, only a limited number of studies have considered the challenge of defining efficient tree proposals. A thorough analysis of standard tree proposals was conducted by Lakner et al. (2008) that provided insight on their performance. Following this study, the concept of guided tree proposals was presented by Höhna and Drummond (2012). This important contribution suggested using scores (e.g., conditional clade probability or posterior probability) to bias the selection of tree alterations toward the most promising among the set of moves proposed by a traditional tree proposal. However, the practicality of the resulting guided proposals remains limited due to the additional incurring computational burden. This concept was nonetheless implemented in MrBayes under the form of parsimony-guided proposals (Ronquist et al. 2012).

Building efficient proposals for continuous parameters has been the subject of numerous studies in the field of computational statistics (e.g., Gelman et al. 1996). These studies have led to the development of adaptive proposals that are self-tuned during an MCMC run to propose moves tailored specifically for the posterior distribution currently estimated (Haario et al. 2001, 2005; Roberts and Rosenthal 2009). The field of computational phylogenetics has employed these approaches to improve the sampling efficiency of continuous parameters by designing novel adaptive proposals (Thawornwattana et al. 2017), developing multivariate proposals that exploit the correlation existing between parameters (Baele et al. 2017; Meyer et al. 2017), or by considering the estimation of distributions approximating the probability distributions of specific parameters (i.e., branch lengths) to independently generate new parameters values (Aberer et al. 2015; Claywell et al. 2018). Most of the software for Bayesian inference of phylogeny take advantage of these methods for continuous parameters (e.g., Ronquist et al. 2012; Aberer et al. 2014; Höhna et al. 2016; Baele et al. 2017). However, as no theory is readily available from the field of computational statistics regarding the sampling of tree topologies, none of these implementations use adaptive proposals for tree topologies.

Software for Bayesian inference of phylogenies continue to mostly rely on tree proposals that naively explore the posterior distribution, as other alternatives are computationally expensive or impracticable. The performance of these proposals hinders our ability to infer large phylogenies and to consider more complex and realistic evolutionary models. In this study, I present the theoretic foundations for the development of adaptive tree proposals and use them to develop three adaptive proposals: two adaptive variants of commonly used proposals and a fully-adaptive proposal based on a novel design philosophy. I investigate the computational complexity of these proposals and define a practical performance metric that enables us to assess the efficiency of each proposal within a mixture. Using this metric, the practical performance of these proposals is then studied on simulated and empirical datasets and compared to the performance of traditional and parsimony-guided tree proposals.

## Materials and methods

### Phylogenetic Tree Proposals

I consider the problem of developing efficient proposal kernels for unrooted tree topologies to conduct Bayesian inference of phylogeny. This type of analysis requires the estimation of the posterior probability distribution:

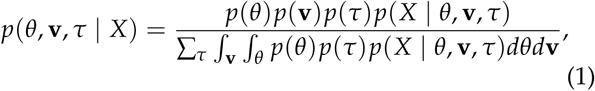

with *θ* being the parameters of the evolutionary model, *τ* the unrooted tree topology, **v** the branch lengths and *X* the alignment. Generally, the posterior distribution is estimated using the Metropolis-Hastings algorithm in which new parameters values are generated by a proposal kernel and accepted with probability

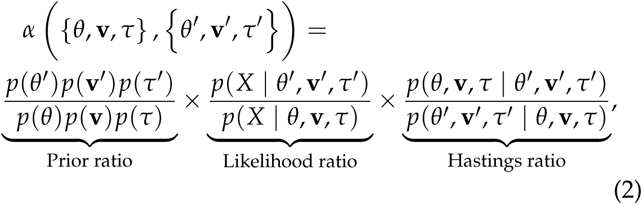

with *p*(*θ*′, **v**′, *τ*′ | *θ*, **v**, *τ*) defining the probability with which the kernel proposes parameters (*θ*′, **v**′, *τ*′) given (*θ*, **v**, *τ*). In this study, I focus on proposal kernels modifying uniquely the tree topology (i.e., *θ* = *θ*′ and **v** = **v**′) and on deriving their Hastings ratios having the form *p*(*τ*|*τ*′)/*p*(*τ*′|*τ*).

Common proposals employed for the inference of unrooted tree topologies are the stochastic Nearest Neighbor Interchange (stNNI), extending Subtree Pruning and Regrafting (eSPR; Swofford et al. 1996) and extending Tree Bisection and Reconnection (eTBR; Huelsenbeck et al. 2008). These proposals naively explore the space of tree topology by arbitrarily altering the current tree *τ* using subtree swapping or pruning operations. For instance, the stNNI proposal interchanges two subtrees separated by an internal branch, while the eSPR proposal prunes a subtree, moves it along a contiguous set of branches (a path) and finally regrafts it on the last branch of the path. Choices of branches, subtrees or paths are made randomly during these proposals and therefore frequently result in tree alterations that may be detrimental (e.g., removing a branch strongly supported by the data).

To improve the quality of the generated moves, adaptive proposals for continuous parameters use summary statistics of the posterior distribution, learned during an MCMC run, to tune the proposal mechanism (Roberts and Rosenthal 2009). Using these summary statistics, parameters of a proposal kernel (e.g., the scale of the random-walk) are adapted to target an optimal acceptance rate. While the specifics of such adaptive proposals are not directly applicable to tree topology proposals, the concept of using summary statistics of the posterior distribution can still be exploited to our advantage. In this study, I consider using the estimated marginal posterior probabilities of splits, or split frequencies, to construct adaptive tree proposals. This strategy is based on two components: the estimation of the split frequencies and the design of adaptive proposals exploiting these estimates.

### Split Frequencies

Each branch of a phylogenetic tree represents a unique bipartition of the set of taxa in the alignment. These bipartitions, better known as splits, are a useful tool to summarize the posterior distribution of trees. Using samples collected during an MCMC run, the marginal posterior probability of a split can be estimated by observing the frequency with which a given split occurs within the sampled tree topologies. Split frequencies therefore provide inexpensive estimates of the support for each specific bipartition but fail to fully summarize the tree distribution by disregarding the joint distribution of splits.

Adaptive proposals require the split frequencies to be learned and made available during MCMC runs. This procedure of estimating split frequencies during a MCMC run is commonly conducted to diagnose the convergence of a set of MCMC runs (Ronquist et al. 2012). As in the post-MCMC estimation of the marginal split frequencies, this procedure is conducted by counting the occurrence of splits within the sampled tree topologies and normalizing them by the number of observed samples. This procedure guarantees that estimates of the split frequencies converge to the true posterior distribution when the MCMC algorithm is run for an infinite amount of time. However, in practice, phylogenetic inferences are not run long enough to ensure the robustness and accuracy of the estimated split frequencies. To tackle this problem, I develop a heuristic algorithm to learn the split frequencies while overcoming several potential issues (SI, Learning split frequency).

The first issue results from the bias induced by the starting parameters of the MCMC. The earliest phase of an MCMC run generally samples trees unrepresentative of the high probability region of the posterior distribution until equilibrium is reached. Therefore, in a post-processing context, split frequencies are evaluated using samples remaining after the removal of the samples collected during the burnin phase. The duration of the burnin phase is however undetermined during an MCMC run and cannot be removed leading the split frequencies to be biased by these early samples. The second issue results from the high volatility and oscillatory behavior of the estimated split frequency during the earliest phase, even when the equilibrium is reached. This oscillatory behavior depends on whether a split is present or absent in the sampled tree topology. Adaptive proposals constructed with these fluctuating estimates could induce unwanted dependencies between the proposal probabilities and the presence or absence of a split (e.g. non-reversibility of the proposals), and impact the correctness of the MCMC algorithm.

For these reasons, the heuristic learning algorithm averages the split frequencies over a fixed number of samples to reduce the volatility of the estimates, uses a relaxation mechanism that progressively reduces the impact of the earliest samples observed, and detects the convergence of the learning process when the amount of variation in split frequencies stabilizes. Upon convergence of the estimated split frequencies, the learning process is terminated to ensure that the proposals preserve the ergodicity of the MCMC algorithm.

Adaptive proposals strongly rely on the estimated split frequencies and, while this heuristic algorithm improves the robustness and accuracy of these estimates, it does not ensure a foolproof estimation procedure. Therefore, to further improve the robustness of adaptive proposals, these proposal mechanisms include a stochastic component *ϵ* to ensure that all trees remain accessible regardless of the accuracy or correctness of the split frequency estimates (SI, Stochastic component).

### Adaptive Tree Proposals

Adaptive tree proposals rely on split frequencies to define regions of the tree that are weakly or strongly supported (i.e., having low or high split frequencies). These regions are used to define moves maintaining highly supported regions of the topology while proposing modifications to regions having weak support. For instance, applying this concept to the stNNI proposal could reduce the frequency of subtree swaps removing a split with strong support while increasing the frequency of swaps acting on a split with weak support.

While this concept can be applied to build adaptive versions of naive proposals (e.g., stNNI or eSPR), two limitations to this approach must be accounted for First, an adaptive version of a proposal always has a larger computational overhead compared to its naive version. To be efficient, an adaptive version must therefore remain computationally inexpensive with respect to the likelihood evaluation induced by a move.

The second key limitation results from a more conceptual consideration: the structure of moves generated by a tree proposal. Naive tree proposals produce moves with a specific structure. For instance, the stNNI proposal interchanges two subtrees separated by a branch while the eTBR proposal generates moves that remove a branch (bisection) and consider a different re-connection of the two resulting subtrees. This design philosophy was fundamental to broaden the range of strategies used to explore tree distributions because each dataset benefits differently from each specific structure of move (Lakner et al. 2008). Ironically, imposing specific structure to moves may prove a hindrance for adaptive tree proposals for the exact same reasons: specific structures of moves do not benefit all datasets.

I propose here a different design philosophy were adaptive tree proposals generate moves having an *adaptive* structure. The information contained in split frequencies represents more than a guiding mechanism for naive proposals. It identifies weakly and strongly supported regions of a tree and their respective structure. These structures provide insight on the structure of moves (e.g., subtrees swap, or pruning and regrafting) that would be appropriate to alter a tree topology. By using the split frequencies to define the structure of moves, adaptive tree proposals have the potential to propose tree alterations specifically tailored to fit the posterior distribution sampled.

In this study, we consider both design philosophies of designing adaptive proposals: proposals producing specific type of moves and proposals using adaptive structure of moves. I first define two adaptive variants of existing naive proposals (stNNI and eSPR) that produce specific type of moves. Then, I present a novel proposal that uses the split frequencies to define moves having an appropriate structure.

#### Mathematical notation

I use the following mathematical notation to describe adaptive proposals (summarized in Table 1): a tree topology *τ* is defined by a set of vertices, *V*, and a set of edges, or branches, *E*. The set of edges, *I*(*E*), identifies the set of internal edges. Each edge *e*_*i*_ identifies a split *s*_*j*_ = *S*(*e*_*i*_) whose frequency is estimated by the function *π*(*s*_*j*_). A split *s*_*j*_ identifies a unique bipartition of the set of taxa and can therefore be identified differently in several tree topologies. For instance, two edges in different trees (e.g., *e*_*i*_ in *τ* and *e*_*j*_ in *τ*′) can identify the same split *s*_*j*_.

**Table 1:**
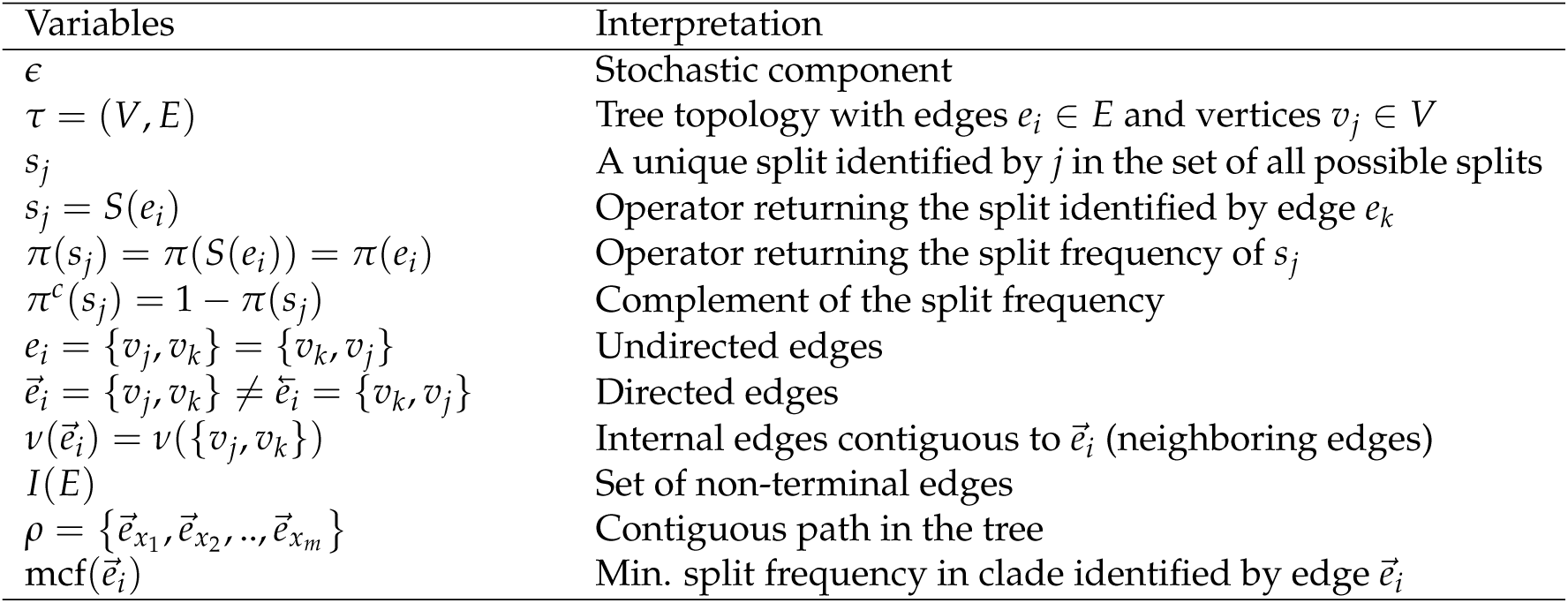
Mathematical notation.

I use a flexible definition of splits in the sense that a split can be used to identify a bipartition and also to build new partitions. For instance, if a directed edge 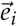 is considered in the context of a move, then the split 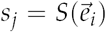 identifies the taxa in the clade subtended by edge 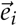. Using this definition, a new split can then be constructed when a clade is moved by considering the union of two splits: for instance, 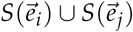 would identify a bipartition segregating the taxa in the clade subtending edges 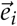 and 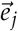 from all the other taxa.

Edges contiguous to edge 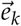 are identified by the operator 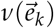 that returns the next edges according to the direction of 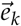. As most of the adaptive proposals considered act on internal edges, the *v*(·) operator only returns edges contained in *I*(*E*). Regions of the tree topology are identified by contiguous paths *ρ* composed of a set of contiguous undirected or directed edges (e.g., 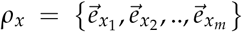). Lastly, the operator mcf 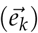 returns the smallest split frequency of the internal edges existing in the clade subtended by edge 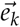.

### Adaptive stNNI

The adaptive stNNI (A-stNNI) proposal uses the split frequencies to guide the selection of the central edge *e*_*r*_ (Fig. 1) whose split *s*_*r*_ = *S*(*e*_*r*_) will be altered by the interchange of two subtrees located on each extremity of *e*_*r*_. The choice of the interchange to apply, among the two possible outcomes, is also guided by the split frequencies.

**Figure 1:**
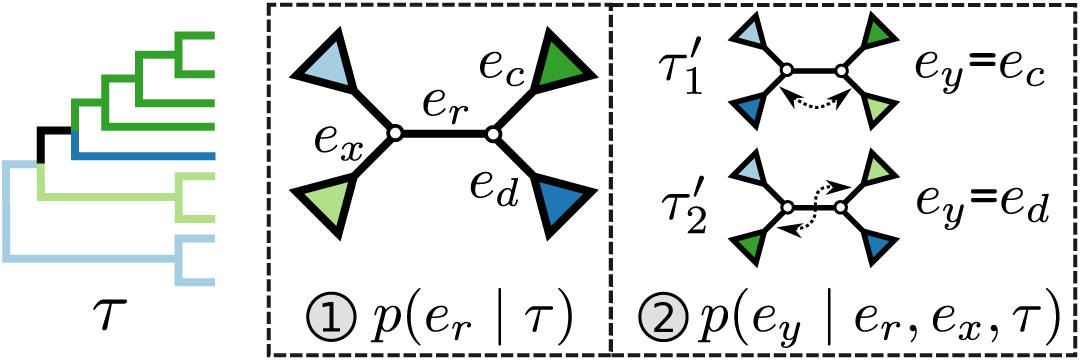
Steps of the A-stNNI proposal. In the first step edge *e*_*r*_ is selected according to Eq. (3). Then, the two subtrees to swap are selected according to Eq. (4).

The selection of the central edge *e*_*r*_ is biased toward edges identifying weakly supported splits. The central edge *e*_*r*_ is therefore selected with probability

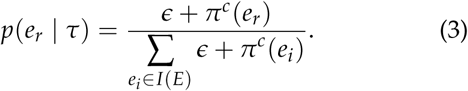

The new split *s*_*u*_ replacing *s*_*r*_ is determined by one of the two possible outcomes of subtrees interchange. The edge 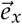 subtending the first subtree to swap is arbitrarily chosen among the four edges contiguous to edge *e*_*r*_ (i.e., with probability *p*(*e*_*x*_|*e*_*r*_, *τ*) = 1/4). The second sub-tree is selected among the two subtree on the opposite side of *e*_*r*_ that are identified by edges *e*_*c*_ and *e*_*d*_, respectively (Fig. 1). The new split *s*_*u*_ segregates the taxa identified by the edge *e*_*x*_ and either edge *e*_*c*_ or *e*_*d*_ from the others, resulting in split *S*(*e*_*x*_) ∪*S*(*e*_*c*_) or *S*(*e*_*x*_) ∪ *S*(*e*_*d*_) respectively. To favor the interchange leading to the tree with the strongest support, the edge *e*_*y*_ identifying the second subtree is therefore selected with probability proportional to *π*(*s*_*u*_):

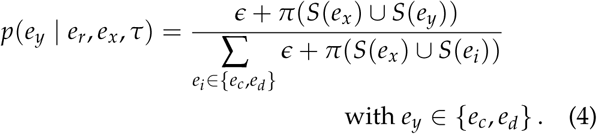

An A-stNNI move is identified by the triplet of edges (*e*_*r*_, *e*_*x*_, *e*_*y*_) and is proposed according to probability

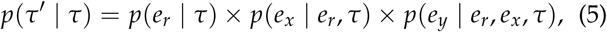

and leads to the new tree topology *τ*′. The reverse move happens with probability

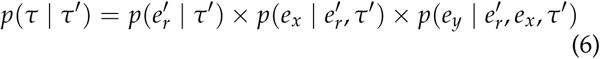

with 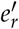 identifying the edge of the new split *s*_*u*_ (i.e., central edge) as in *τ*, and edges *e*_*x*_ and *e*_*y*_ identifies the same splits in *τ* and *τ*′. The Hastings ratio required to evaluate the acceptance probability of a A-stNNI move (Eq. (2)) is the ratio of Eqs. (6) and (5).

### Adaptive 2-edges SPR

The adaptive 2-edges SPR (A-2SPR) is an adaptive version of the eSPR move. Similarly to the eSPR proposal, this adaptive proposal prunes a subtree, moves it along a path made of consecutive edges, and regrafts it (Fig. 2). The length of the path is however limited to exactly two edges, resulting in the alteration of two splits exactly. Conversely to the usual eSPR strategy, the A-2SPR first selects the path along which the subtree will be moved and then consider all the possible pruning and regrafting moves along this path. The A-2SPR can be seen as a natural extension of the A-stNNI to two edges since the path is selected to target region of the tree with weak support, while the move is selected to favor the resulting tree having the strongest support.

**Figure 2:**
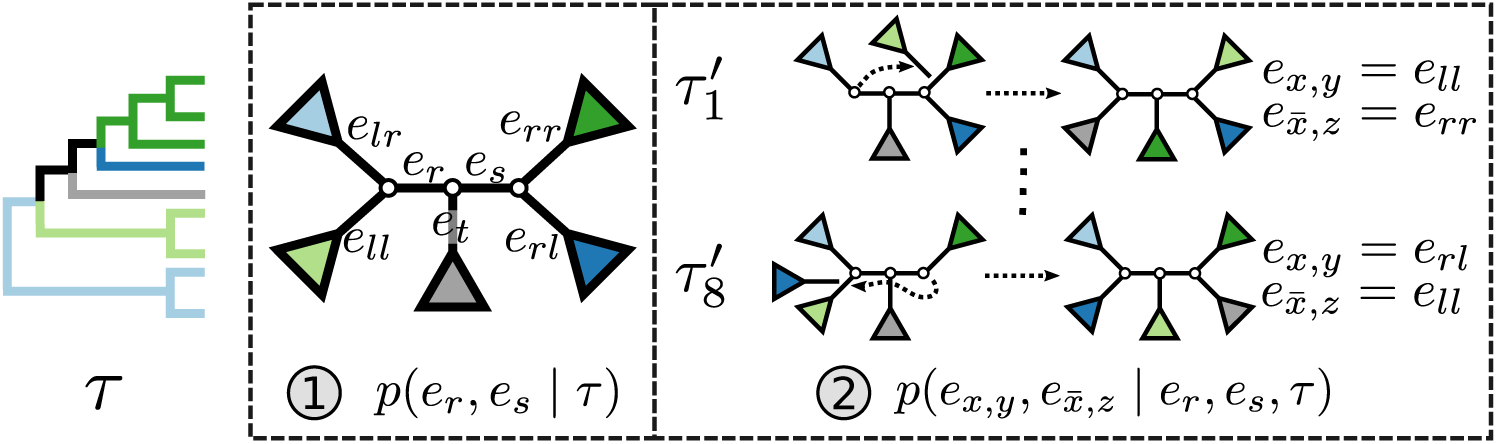
Steps of the A-2SPR proposal. In the first step the edges (*e*_*r*_, *e*_*s*_), along which a subtree will be moved, are selected according to Eq. (7). Then, a move among the 8 possible pruning and regrafting locations along edges (*e*_*r*_, *e*_*s*_) is selected according to Eq. (8).

In the first step, I select a pair of contiguous edges (*e*_*r*_, *e*_*s*_) with probability inversely proportional to the product of their estimated marginal split frequencies, as defined by,

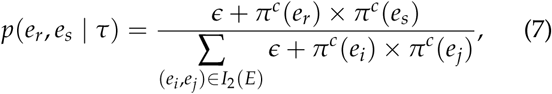

where *I*_2_(*E*) identifies the set containing all pairs of contiguous internal edges in *τ*. The pair (*e*_*r*_, *e*_*s*_) has the four edges *e*_*a*_, *e*_*b*_, *e*_*c*_ and *e*_*d*_ at its extremities and the edge *e*_*t*_ in its center, which is the edge sharing a vertex with both edges *e*_*r*_ and *e*_*s*_.

The second step enumerates all the possible moves across edges (*e*_*r*_, *e*_*s*_) for subtrees subtended by edges *e*_*a*_, *e*_*b*_, *e*_*c*_ and *e*_*d*_. Each of those edges can be regrafted two ways after moving along edges (*e*_*r*_, *e*_*s*_). For instance, assuming that *e*_*a*_ and *e*_*b*_ are adjacent, the subtree identified by edge *e*_*a*_ could be pruned and then regrafted on edges *e*_*c*_ or *e*_*d*_. Assuming that edge *e*_*c*_ is selected as the regrafting point, the split *s*_*r*_ = *S*(*e*_*r*_) and *s*_*s*_ = *S*(*e*_*s*_) would be removed and would be replaced by splits *s*_*p*_ and *s*_*q*_. The first split would separate taxa in subtrees identified by edges *e*_*b*_ and *e*_*t*_ from the others (i.e., *s*_*p*_ = *S*(*e*_*b*_) ∪ *S*(*e*_*t*_) = *S*(*e*_*a*_) ∪ *S*(*e*_*c*_) ∪ *S*(*e*_*d*_)), while the second would separate taxa in the clade identified by edges *e*_*a*_ and *e*_*c*_ from the others (i.e., *s*_*q*_ = *S*(*e*_*a*_) ∪*S*(*e*_*c*_)).

For mathematical convenience, edges *e*_*a*_, *e*_*b*_, *e*_*c*_ and *e*_*d*_ can be relabeled using their relative position using notation 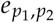 (Fig. 2). Indexes *p*_*i*_*∈*{*r, l*} defines whether the edges are at the right (*p*_1_ = *r*) or left (*p*_1_ = *l*) extremity of edges (*e*_*r*_, *e*_*s*_) and whether the edges are the right or left one relatively to each other (*p*_2_). Using this notation for the edge *e*_*x,y*_ identifying the subtree to move at extremity *x* of edges (*e*_*r*_, *e*_*s*_) and the edge 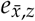 identifying the regrafting point at the opposite extremity 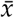 (e.g.. 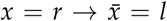), the probability of selecting a move that favors the strongest supported tree is defined as

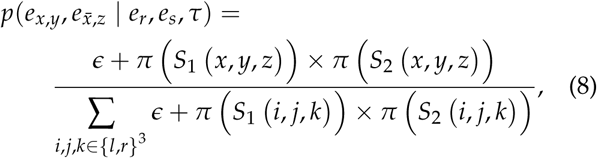

with

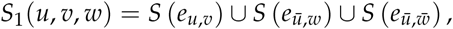

and

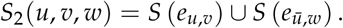

The move identified by the quadruplet of edges 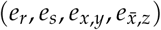 is proposed according to probability,

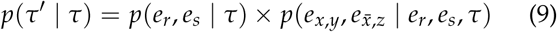

and leads to the new tree topology *τ*′. The reverse moves happens with probability,

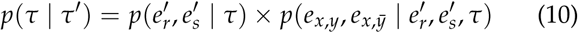

where edges 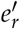 and 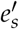 identify the new splits. Edge *e*_*x,y*_ identifies the edge originally pruned and 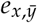 its neighbor in *τ* that subtend to the same subtrees in *τ* and *τ*′. The ratio of Eqs. (10) and (9) defines the Hastings ratio for the A-2SPR proposal that is used to evaluate the acceptance probability of a move (Eq. (2)).

This strategy can be generalized to build adaptive N-edges SPR moves. However, two pitfalls are inherent to this approach. First, using a proposal affecting a fixed number of edges is inconvenient and can result in nonergodic proposals when used on their own. Second, the proposals efficiency would suffer from the increasing computational cost. The computational complexity of the enumeration of all *N*-edges paths grows as *O*(2^*N*^), while the one of estimating the move probabilities grows as *O*(*N*) (SI, Computational complexity).

### Adaptive Path Building and Jolting Proposal

While the A-stNNI and A-2SPR proposals are based on a design philosophy targeting the construction of specific type of moves, the adaptive path building and jolting proposal (A-PBJ) embraces the design philosophy of using the split frequencies to adaptively define the proposals structure. This approach is achieved by using the split frequencies to identify two types of structures within a tree topology: weakly and strongly supported regions. A proposal designed under this philosophy should strive to maintain the regions with strong support while proposing alteration to ones with weak support. Moves resulting from this strategy are specifically tailored to fit the posterior distribution of trees at hand.

The A-PBJ proposal implements this novel design philosophy by using contiguous paths within a tree to identify weakly and strongly supported regions: the unstable and stable paths, respectively. This proposal begins by selecting a stable path acting as the backbone of the move. Unstable paths are then constructed at both extremities of the backbone and are subject to eSPR-type moves. In two separate steps, each edge at the extremities of the stable path are pruned, moved along their relative unstable path and then regrafted (Fig. 3). The stable path and its splits remain unaltered by this move, while the splits of both unstable paths are replaced by new ones. Depending on the paths identified, moves produced by the A-PBJ proposal include (but not exclusively) moves produced by the stNNI, eSPR and eTBR proposals.

**Figure 3:**
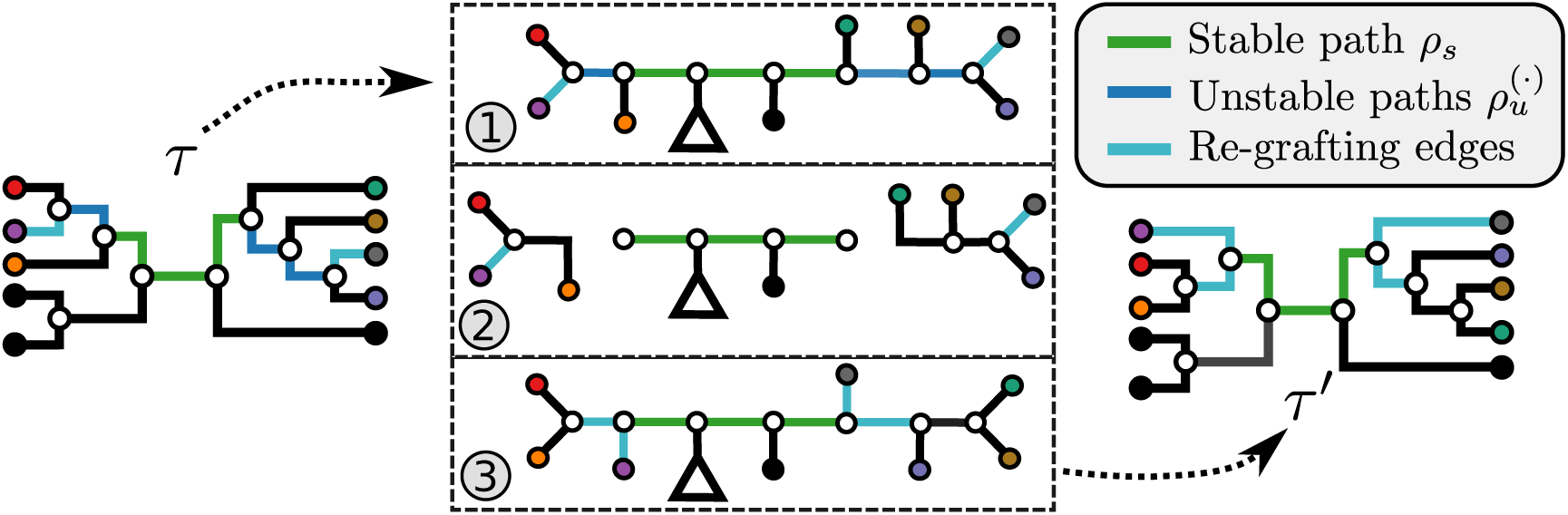
Example of an A-PBJ proposal. The two eSPR-type moves defined by the stable path *ρ*_*s*_ (Fig. 4) and unstable paths 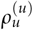 and 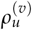 (Fig. 5) are applied on the example tree *τ*, resulting in tree *τ*^*t*^. The probability of an A-PBJ move is the joint probability of building the three paths (Eqs. (11) and (12)).

The path building strategies are key to the efficiency and reliability of the A-PBJ proposal. The stable paths must capture sets of splits having high frequencies that would benefit from alterations at their extremities; constructing stable paths starting and ending at terminal nodes would not enable any moves. The unstable paths must capture set of low-frequencies split of variable size and, in this sense, act as a generalization of the mechanisms previously employed in the A-stNNI and A-2SPR. This generalization must however avoid the expensive enumeration of all possible N-edges paths to remain computationally competitive.

The strategies used to construct such paths and their resulting probabilities are defined in the following sections. The construction of a stable path *ρ*_*s*_ with probability *p*(*ρ*_*s*_|*τ*) is illustrated in Fig. 4. The unstable paths 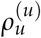 and 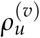 are built at each extremities of the stable path *ρ*_*s*_ identified by edges 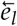 and 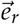 The constructions of those unstable paths have probabilities 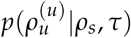 and 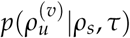, respectively, and are illustrated in Fig. 5.

**Figure 4:**
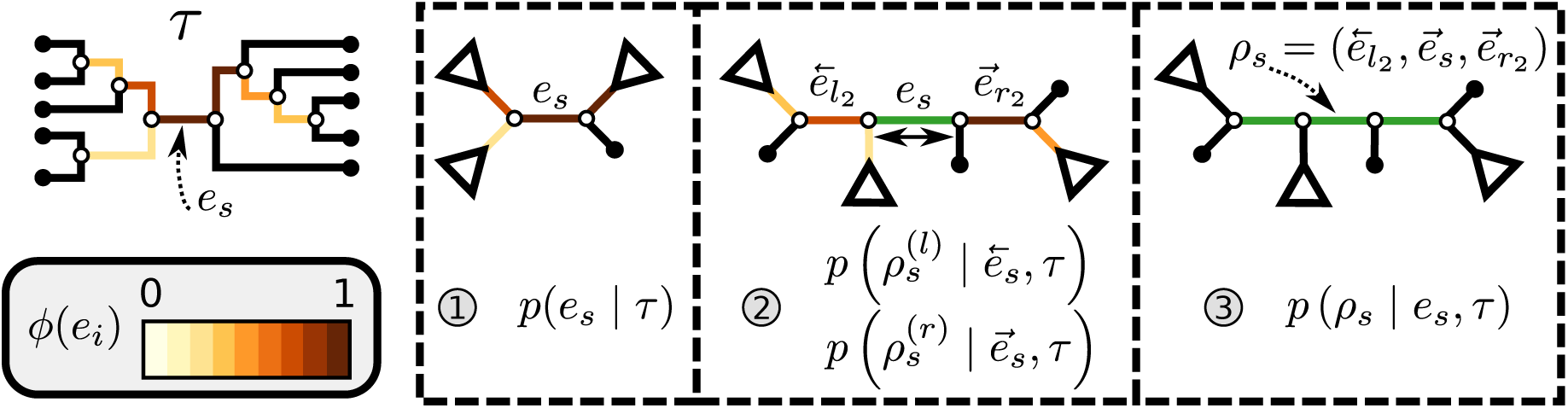
Construction of a stable path *ρ*_*s*_. In the first step the edge *e*_*s*_ is selected according to Eq. (13). Then, the path is extended on both side of edge *e*_*s*_ according to Eqs. (14-16). Finally, both extensions are concatenated forming path *ρ*_*s*_ (Eqs. 17-19)).

**Figure 5:**
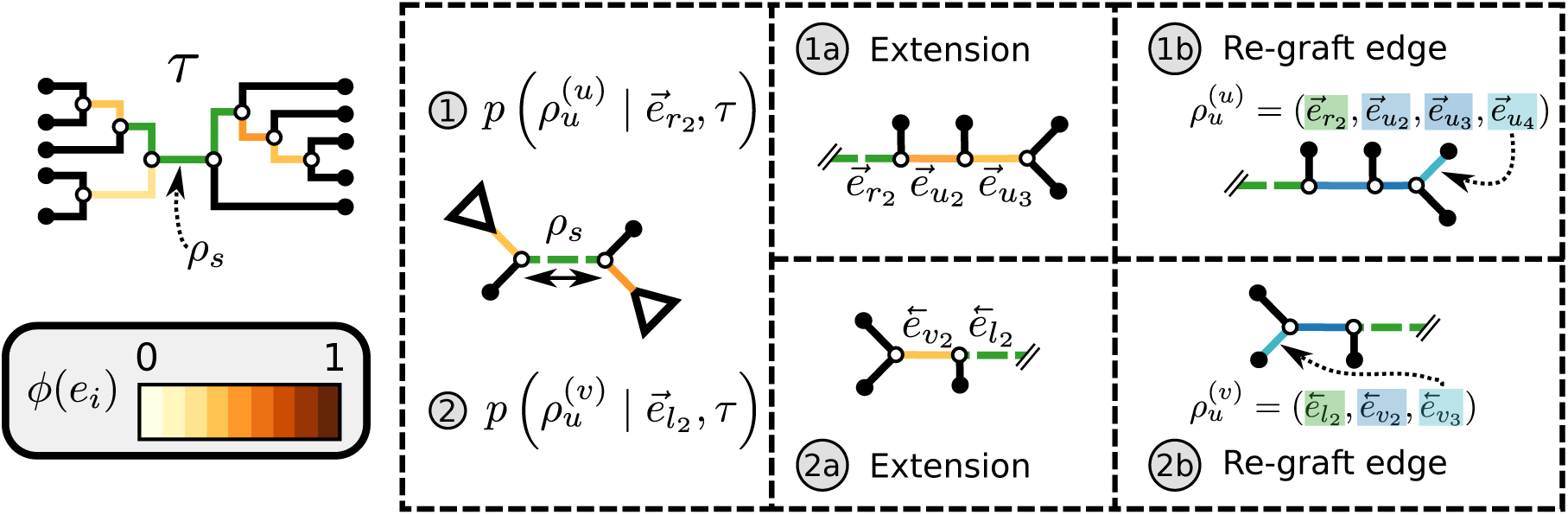
Construction of the unstable paths 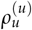 and 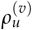 at the extremities of *ρ*_*s*_ (Fig. 4). In the two separate building phase (1 and 2), the paths are constructed by extension (1a and 2a) according to Eqs. ((20)-(22)). Then, the last edges identifying the re-graft points (1b and 2b) are selected according to Eq. (23). The probability of building paths 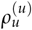 and 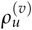 is the joint probability of the extensions and re-graft selections (Eq. (24)).

The probability of an A-PBJ move is defined using the joint probability of building its composing paths and is given as

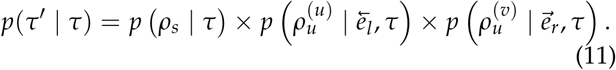

The reverse move happens with probability

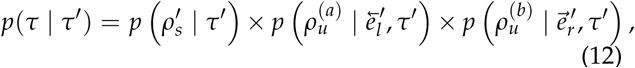

where 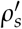 identifies the path including edges identifying the same splits as *ρ*_*s*_. Paths 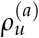 and 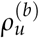 are inverse versions of path 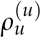 and 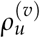, respectively, that identify the reverse eSPR-type moves.

The ratio of Eqs. (12) and (11) defines the Hastings ratio for the A-PBJ proposal required to evaluate the acceptance proposal of a move (Eq. (2)). The different terms involved in Eqs. (11) and (12) are detailed in the next sections.

#### Stable path

The construction of a stable path *ρ*_*s*_ consists first in the selection of an edge with probability

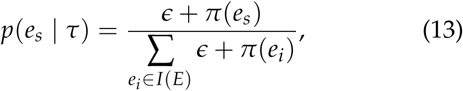

that favors the selection of internal edges identifying high-frequency splits. The path *ρ*_*s*_ is then built by stepwise extension of its extremities. Starting from edge *e*_*s*_ the path is extended in both directions, namely the 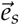 and 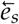 directions, respectively (Fig. 4). After including edge *e*_*s*_ in a partial path (e.g., 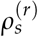), the extension mechanism iterates over two steps: first, the termination condition of the extension, and second (if not terminated), the extension continues by choosing the new edge to add to the path.

Assuming an initial direction 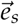 and the initial partial path 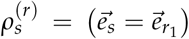 with the first iteration (*i* = 1) begins by controlling the termination condition that occurs with probability

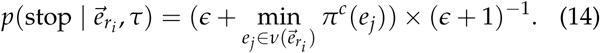

This probability favors the termination of the extension mechanism whenever a split identified by a neighboring internal edge 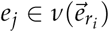 could benefit from being altered.

If the termination condition is not met (which happens with probability 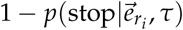), the path continues its extension by selecting the next edge to add, if any. Nonterminal edges identifying a split with high frequency and leading to a clade containing low-frequency structures represent good candidates for extension and are selected with probability

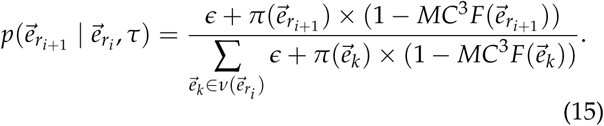

The selected edge 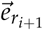 is then added to the path (i.e., 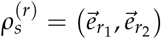) and a new iteration begins (*i* = *i* + 1).

This process continues until the termination event occurs or until the path reaches an endpoint (i.e., 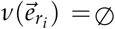). The probability of having extended the partial stable path 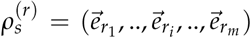 in direction 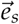 is then given as

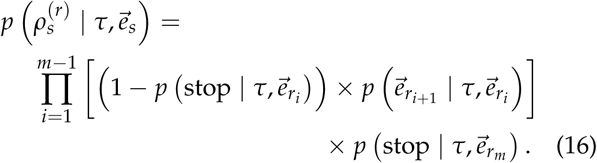

The extension mechanism is repeated in the opposite direction (i.e., 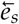) resulting in partial stable path 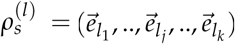 Both partial paths 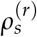 and 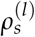 are then concatenated to form the stable path

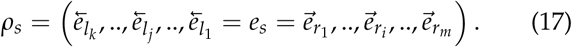

The construction of this path is conditional on the selection of edge *e*_*s*_ as starting point and is built with probability

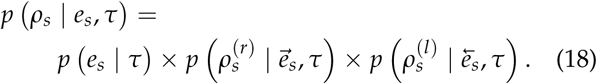

Given that this exact path may be built starting from any edge *e*_*i*_ *∈ ρ*_*s*_, the probability of building path *ρ*_*s*_ must be marginalized over all potential starting edge and is defined as

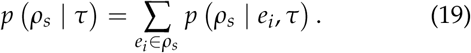

#### Unstable paths

The mechanism used to build unstable paths consists of extension phases starting from each of the extremities of the stable path *ρ*_*s*_, each identified by edges 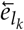 and 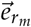, respectively. An unstable path fully identifies an eSPR-type move (Fig. 5). For instance, the unstable path 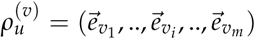 starting from edge 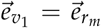 includes the edge to prune 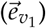 that identifies the moving clade *C*, the edge that identifies the direction of the move 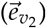, the edges traversed by clade *C* and the regrafting point of clade *C* 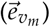. As in the extension of the stable path, an unstable path 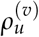 is built by iterating over two steps: the extension termination and edge selection steps.

The extension phase is preferably terminated when the unstable path would risk begin extended by an edge identifying a split with a high-frequency in the next step. The termination event occurs then with probability

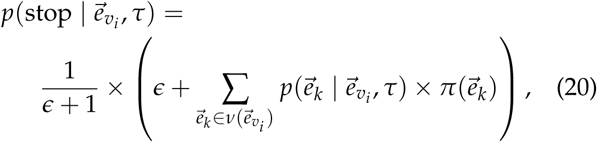

where 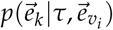 identifies the probability of extending path 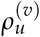 with edge 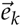 (Eqs. (21-23)).

If the termination does not occur, the next edge is selected according to probabilities defined by its role in the eSPR move (i.e., direction, traversed or regrafting edge). The first edge selected identifies the direction along which clade *C* will move and is selected with probability

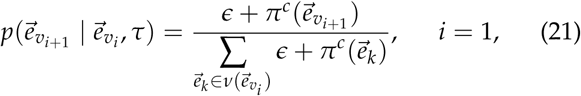

that favors the removal of edges identifying low-frequency splits.

Each edge traversed by clade *C* is then selected with probability

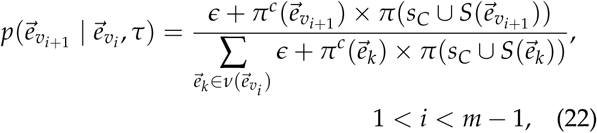

where *s*_*C*_ identifies the split containing the taxa of clade *C*. This probability accounts for the removal of the current split 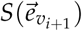 and the addition of the new split 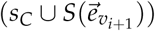.

The last edge, selected after the termination of the extension phase, identifies the regrafting edge for clade *C* and is selected with probability

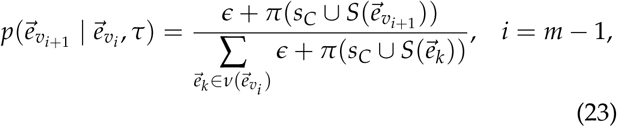

that is, proportionally to the frequency of the last split added.

Building an unstable path 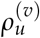 is the outcome of the selection of the direction, the extension of the traversed path and the choice of the regrafting edge. The probability of building an unstable path is therefore defined as,

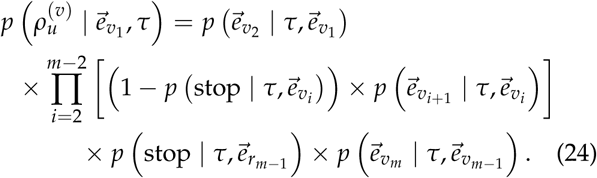

The probability of an A-PBJ move (Eqs (11) and (11)) is defined as the joint probability of building a stable path (Eq. 19) and the two unstable paths starting at its extremities (Eq (24)).

### Parsimony-Guided stNNI and eSPR

I implemented Parsimony-guided stNNI and eSPR proposals based on the concepts presented in (Höhna and Drummond 2012) to compare adaptive proposals with other strategies for guiding tree proposals (SI, Parsimony score transformation). Two different strategies were considered: exhaustive guided stNNI (G-stNNI) and guided N-edges eSPR (G-NSPR) proposals. The mechanism of these proposals consists of defining a set of potential moves and drawing one of them proportionally to the parsimony score of the resulting trees. Each proposal differs in the strategy used to build the set of moves. The G-stNNI proposal enumerates all possible stNNI moves for the current tree *τ*, while the G-NSPR proposal randomly choose sa subtree to prune, then enumerates all eSPR moves altering at most N-edges. Using *N* = 1 has a similar effect to a guided stNNI that would randomly choose the central edge and then use the parsimony score to guide the subtree interchange.

### Theoretical Computational Complexity of Proposals

I defined the theoretical computational complexity of the different proposals and compared them to the cost of a partial likelihood (SI, Computational complexity). These complexities and the condition under which a proposal is computationally efficient with respect to a partial likelihood evaluation are summarized in Table 2. This theoretical analysis indicates that parsimony-guided tree proposals have performance improvements limited by their dependencies to the number of sites *m*. Conversely, adaptive tree proposals should have negligible computational cost as long as the number of taxa *n* is smaller than *m*.

**Table 2:**
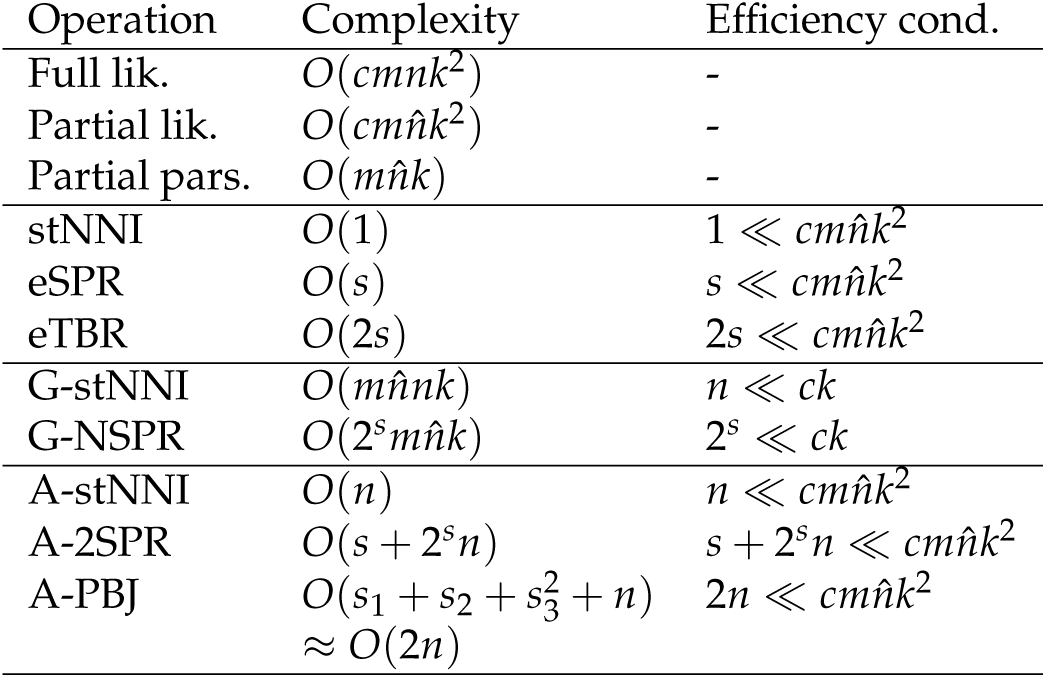
Computational complexity of the proposals and likelihood operations. Alignments have *m* sites for *n* sequences defined over an alphabet of *k* characters (e.g., *k* = 4 for nucleotide sequences). The Gamma model for rate heterogeneity has a number *c* of rate categories (Yang 1994), while a partial likelihood requires 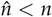 operations instead of *n*. Finally, *s* represents the number of splits altered by a move.

The parsimony-guided proposal G-stNNI could exceed the computational complexity of a partial likelihood evaluation under the condition that the number of taxa *n* exceed the product of the number of symbols *c* (e.g., *c* = 4 for nucleotides) and the number of rate categories *k* under a discrete-Gamma rate model (Yang 1994). Such scenarios would happen even when using moderately complex models as the GTR+Γ substitution model with *k* = 4 rates categories. The G-NSPR proposal seems more competitive as its efficiency condition is reached when 2^*s*^ ≪ *ck*. Even if this condition is not reached, a G-NSPR can be executed at a fraction 2^*s*^/(*ck*) of a partial likelihood evaluation, which represents a reasonable computational overhead for small values of *s*.

While the overhead cost of adaptive proposals is non-negligible compared to naive proposals, their computational overhead remains negligible with respect to partial likelihood evaluations as long as 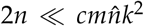, where 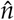 identifies the number of edges included in a partial likelihood evaluation. This efficiency condition is met even for the simple models (e.g., nucleotide substitutions model without rate heterogeneity, *c* = 1, *k* = 4) as long as *n* ≤ *m* (which is a prerequisite to obtain accurate inferences of phylogeny).

### Assessing the Performance of Tree Proposals

Diagnosing the behavior of an MCMC algorithm is generally achieved by monitoring its sampling performance (e.g., effective sample size) on different parameters. This task is particularly difficult when monitoring the performance of MCMC algorithms to estimate the posterior distribution of trees due to the discrete nature of this parameter. Nonetheless, two different characteristics are usually monitored. First, the time to convergence of the MCMC algorithm, that is the number of iterations required until the Markov chain reaches its equilibrium. Once at equilibrium, samples obtained from the Markov chain are representative of the true posterior distribution. The second characteristic is the mixing efficiency, that is the propensity of the MCMC algorithm to mimic the process of directly drawing samples from the true posterior distribution. By exploring the parameter space in a step-wise manner, random-walk MCMC algorithm have the tendency to generate autocorrelated samples that are more predictable than draws from the posterior distribution.

Few robust and practical procedures exist to measure these two characteristics when phylogenetic trees are involved and none are able to separately monitor the behavior of tree proposals within a mixture. In the next sections, I first enumerate the existing procedures and define how I used them, and then I present a novel metric that is able to isolate and assess the performance of tree proposals contained in a mixture of proposals.

#### Existing performance metrics

The standard metric to assess the convergence of MCMC runs estimating the posterior distribution of trees is the average standard deviation of split frequencies (ASDSF). This metric captures the variance of the posterior distribution of split frequencies among several independent MCMC runs. In previous studies, the efficiency of tree proposals was assessed using a convergence threshold based on the ASDSF (Lakner et al. 2008; Höhna and Drummond 2012). After a thorough investigation of this procedure, Whidden and Matsen (2015) suggested that the number of replicates plays a key role in the accuracy of this metric. Instead of the ASDSF, graph metrics such as the mean round trip cover time (MRT) were used to assess the behavior of the MCMC runs. Alas, these metrics could not be applied on diffuse tree distributions and required MCMC runs consisting of an enormous number of samples.

Using large amount of replicates or replicates having enormous numbers of samples to assess the behavior of MCMC runs is impractical when the behavior of many tree proposals are compared on several datasets. Therefore, I used multiple long and independent runs with MrBayes to obtain accurate estimates of the posterior distribution of split frequencies *p*(**s**|*X*). These reference split frequencies were used to assess the convergence of each one of the runs under two scenarios: the overall convergence of the MCMC run (before burnin removal) and convergence after removal of the burnin phase. The first metric provides an indication on the number of iterations required to reach convergence, while the second indicates the efficiency of the proposals at estimating the split frequencies. I defined these convergence criteria as satisfied when the Euclidean distance between the reference *p*(**s**|*X*) and the estimated 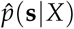 marginal distribution of split frequencies, for all splits *s*_*i*_ with *p*(*s*_*i*_|*X*) *>* 0.1, was lower than a fixed threshold. I defined these thresholds to represent an average error of 2% and 1% per split for the before and after-burnin removal convergence metrics, respectively.

#### Assessing the contribution of moves in a mixture of proposals

The aforementioned convergence metrics capture the behavior of MCMC runs without providing information on the performance of the different tree proposals forming a mixture. With the exception of the acceptance rate of each tree proposal, no metrics are readily available to measure each the relative contribution of each proposal. Furthermore, while the acceptance rate provides some information on the proposal behavior, it fails to capture the amount of trees that the proposal can access or alter: a proposal that would efficiently visit only a small subset of the 95% highest posterior density (HPD) would have a high acceptance rate. Additionally, measuring the convergence metrics on finite MCMC runs using a single tree proposal at a time could result in failure to reach convergence for some proposals. Failure to reach convergence is unrepresentative of the mixing efficiency of tree proposals and could therefore lead us to discard proposals that efficiently sample the posterior distribution of trees once convergence is reached.

I therefore considered an alternative approach to characterize the performance of each tree proposal in a mixture. This strategy considers as reference the expected behavior of an idealized proposal: the joint posterior distribution of parameters *θ*, **v** and trees *τ*. Instead of basing these metrics on a hypothetical sequence of trees produced by this proposal, I investigate the use of summary statistics on the number of iterations between different visits of a given split. A cycle corresponding to the visit of a split begins with its appearance in the trees sampled by the MCMC process. It then includes the number of subsequent iterations when the split is still sampled, plus the number of iterations when the split is not sampled, until its first re-appearance. When trees are drawn directly from the posterior distribution, the frequency at which a split is visited depends entirely on its posterior probability and can be derived by considering the sum of two geometric random variables (Sen and Balakrishnan 1999). These two geometric distributions characterize the number of samples before the disappearance of the split and then the number of iterations before its reappearance in the trees sampled. The probability of the period *k*_*i*_ of a visit-cycle given the split posterior probability *p*(*s*_*i*_|*X*) is given as,

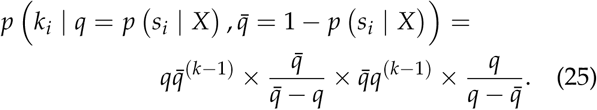

Assuming that accurate estimates of the split frequencies are available (e.g., reference runs), I can summarize the characteristics of the ideal tree proposal with a split-wise approach by considering the expected period *E*(*k*_*i*_) of splits *s*_*i*_ (Fig. 6a and b). This reference behavior can then be compared to the one observed from adaptive proposals, or mixture of proposals, under the condition that their history is tracked by logging the iteration at which they are applied and the resulting effect on the tree sampled (i.e., move accepted or rejected). From this information, the observed average cycle-visit (C-V) period 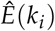 per proposal and per split can be summarized (Fig. 6c to e).

**Figure 6:**
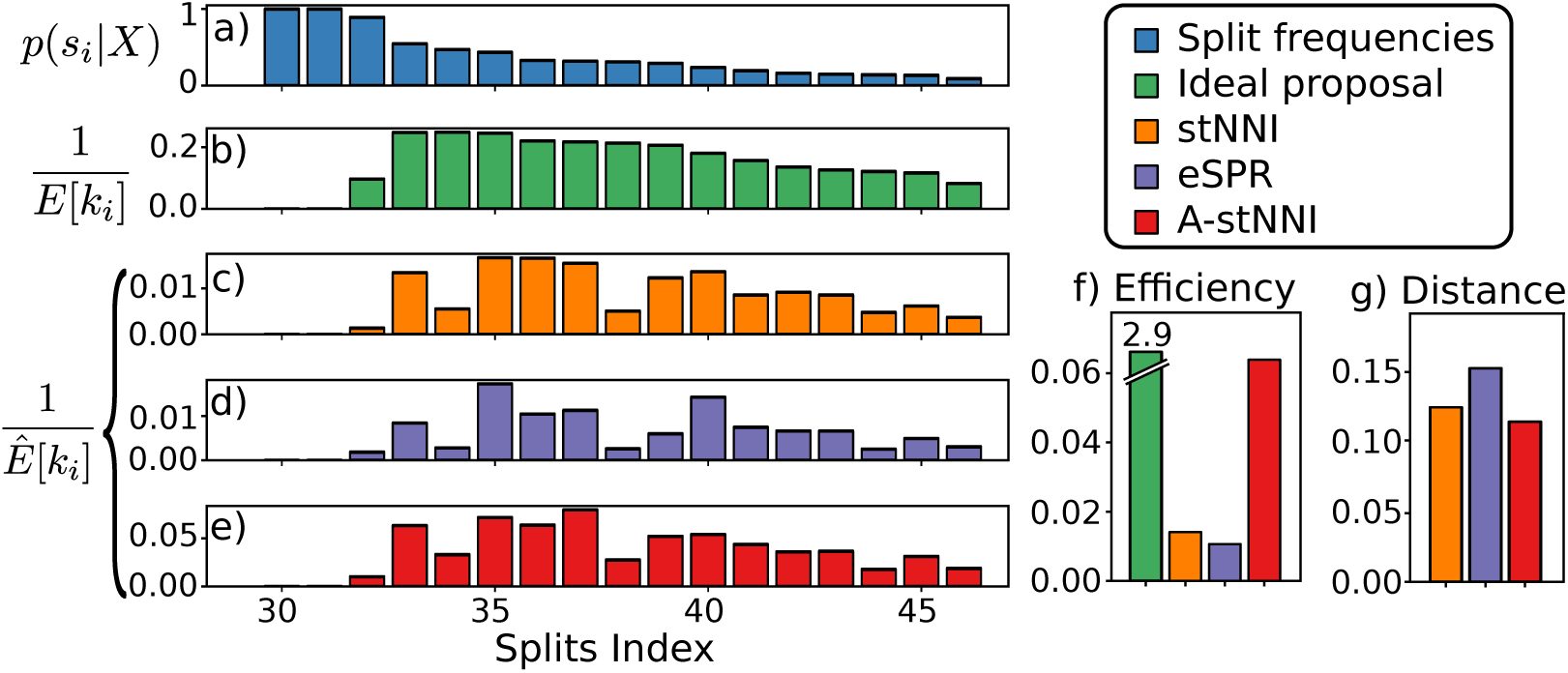
Example for the visit-cycle metrics. Panels a) displays the reference split frequencies. Figure b) shows the ideal C-V frequencies. Figure c) to e) represents the observed C-V frequencies for the stNNI, eSPR and A-stNNI, respectively. Figures f) and g) indicate the resulting C-V efficiency metric and C-V distance from the ideal proposal.

I use the C-V period for each split *s*_*i*_ with *p*(*s*_*i*_|*X*) *>* 0.1, or more exactly their C-V frequency (1/*E*(*k*_*i*_)), to assess the performance of each tree proposal applied within an MCMC run according to two metrics: their efficiency and split coverage. The C-V efficiency is estimated as the sum of the C-V frequencies over all splits and represents the average number of splits visited per moves (Fig. 6f). For proposals altering a single split (i.e., stNNI) this measure is equivalent to the acceptance rate. The second metric is estimated as the difference between the relative C-V frequency of a proposal and the one expected under the ideal proposal. This metric, the C-V, distance is defined as

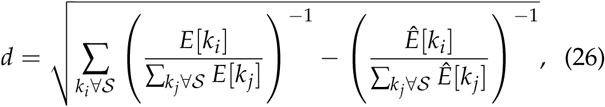

where 𝒮 identifies all cycle-period *k*_*i*_ of splits *s*_*i*_ having a posterior frequency *p*(*s*_*i*_|*X*) *>* 0.1 (Fig. 6g). This distance represents the departure from the expected relative C-V frequencies: proposals visiting mostly a small subset of splits, for instance, would have a behavior distant to the one of the ideal proposal. To simplify the interpretation of the C-V distance in upcoming benchmarks, I will consider the inverse of this metric (*d*^−1^): the C-V accuracy. This transformed metric conveniently increases from 0 to 1 as the proposal behavior approaches the one of the ideal proposal.

## Results

This study is decomposed in two separate experiments. First, I assessed the performance of each proposal separately and validated the new efficiency metrics on datasets simulated with the INDElible software (Fletcher and Yang 2009). Trees were simulated under a birth-death model, while the alignments were simulated under the GTR model (SI, Dataset simulation settings). Then, I assessed the performance of a proposals mixture containing adaptive proposals by comparing it to the traditional mixture of naive proposals. This comparison was conducted using 11 empirical datasets commonly used to evaluate tree proposals (SI, Table 4; Lakner et al. 2008; Höhna and Drummond 2012; Whidden and Matsen 2015).

### Set-up of the Analyses

I analyzed each datasets with MrBayes (Ronquist et al. 2012) to obtain accurate estimates of the tree and split posterior distributions. Simulated and empirical datasets were analyzed under the GTR model and GTR+Γ model with 4 categories, respectively (SI, priors settings). Analyses of at least 50 million iterations with 4 Metropolis-coupled chains were replicated 3 times and used to define the reference split frequencies. Each analysis reached an ASDSF value smaller than 0.005, suggesting that runs converged properly.

The adaptive and parsimony-guided proposals are implemented in the CoevRJ software (Meyer et al. 2019) that I therefore used to assess their performance. Analyses of 10 million iterations with 10 independent chains were conducted for each proposal mixture and dataset with the same models and prior setting used to build the reference split frequencies. Additional runs with 3 Metropolis-coupled chains were conducted for the empirical datasets (using the same settings, iterations and replicas number as their MCMC equivalent).

### Mixtures of Proposals

Five different proposal mixtures were considered for these experiments (SI, Table 3). The reference mixture is the *naive* mixture composed of the stNNI, eSPR and eTBR proposals applied each at equal frequency (as in MrBayes), with a probability *p* = 0.6 of extending a path for eSPR and eTBR. All subsequent mixtures applied stNNI and eSPR proposals at low frequency as a baseline.

**Table 3:** Proposal mixtures and proposal weights. The weights define the relative frequencies at which each proposals are applied.

| Mixture | Proposals (weight) |
| --- | --- |
| *Naive* | stNNI (1), eSPR (1.0), eTBR (1) |
| *Adaptive* | stNNI (1/3), eSPR (1/3),<br>A-stNNI (1), A-2SPR (1), A-PBJ (1) |
| *Guided* | stNNI (1/3), eSPR (1/3),<br>G-stNNI (1), G-1SPR (1), G-2SPR (1) |
| *Mixed* | stNNI (1/3), eSPR (1/3),<br>A-stNNI (1), A-2SPR (1), A-PBJ (1),<br>G-1SPR (1), G-2SPR (1) |
| *Best* | stNNI (1/15), eSPR (1/20),<br>A-stNNI (1), A-2SPR (1/10), A-PBJ (1/5)<br>G-2SPR (1/15) |

**Table 4:**
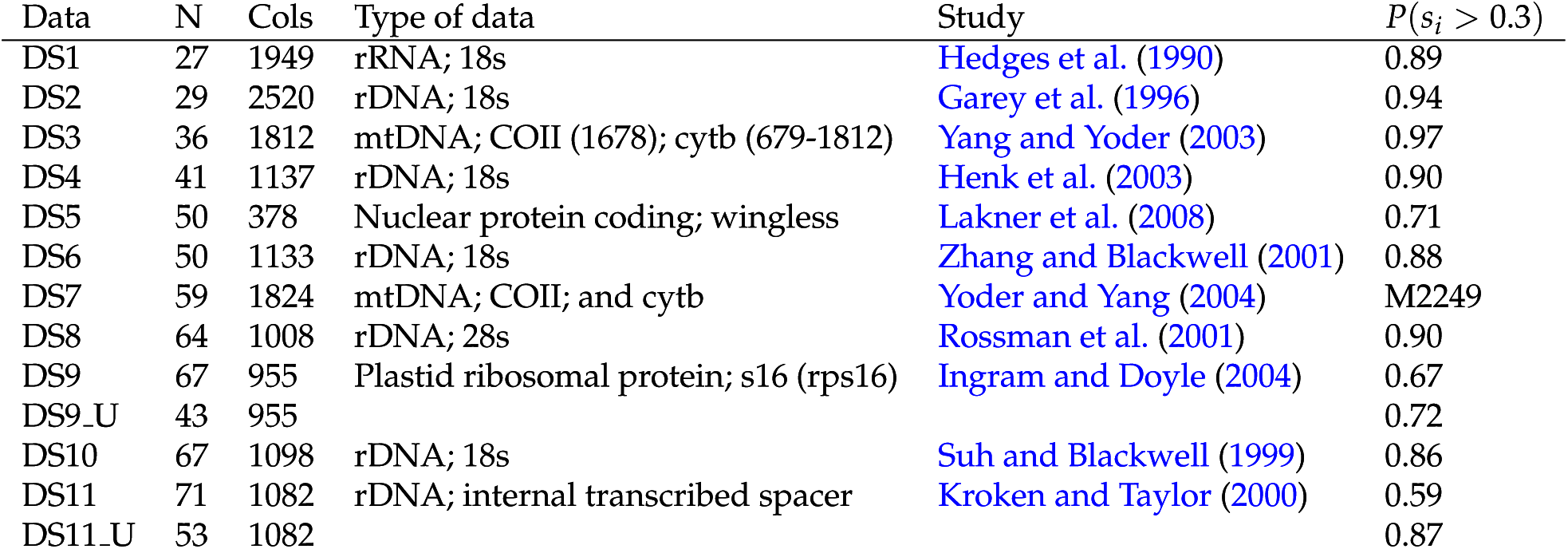
Empirical datasets used and the *information* contained in the split frequencies. Datasets postfixed with the letter *u* have been altered by removing duplicate sequences.

The *adaptive* and *guided* mixtures were composed of all adaptive (A-stNNI, A-2SPR and A-PBJ) and guided (G-stNNI, G-1SPR, G-2SPR) proposals, respectively, each applied at equal frequencies. The *mixed* mixture applied all the adaptive and guided proposals with equal frequencies, with the exception of the G-stNNI that was removed due to its expensive computational cost. Finally, the *best* mixture identified a version of the *mixed* mixture with the proposal weights tuned for performance (based on empirical observations).

### Performance of Proposal Mixtures on Simulated Datasets

#### Proposal performance

On the four simulated data sets, the adaptive proposals achieved the best mixing efficiency according to the C-V efficiency with consistent 2 to 8-fold performance improvements over the stNNI proposal (Fig 7a). These increases in C-V efficiency were mirrored by increased acceptance rates (Fig 7c) while being of slightly lesser extent for proposals modifying multiple splits per move (e.g., A-PBJ). The ability of proposals to visit all splits according to the theoretical expectation, when compared to the stNNI proposal, generally decreased at the exception of the A-stNNI (Fig 7b). This decrease in C-V accuracy was linked with the number of edges altered (e.g., 2-edges moves only for the A-2SPR). Lastly, the measured computational cost for each proposal reflected the theoretical complexity, and particularly affected the G-stNNI proposal with a nearly 10-fold increase in computational cost (Fig 7d).

**Figure 7:**
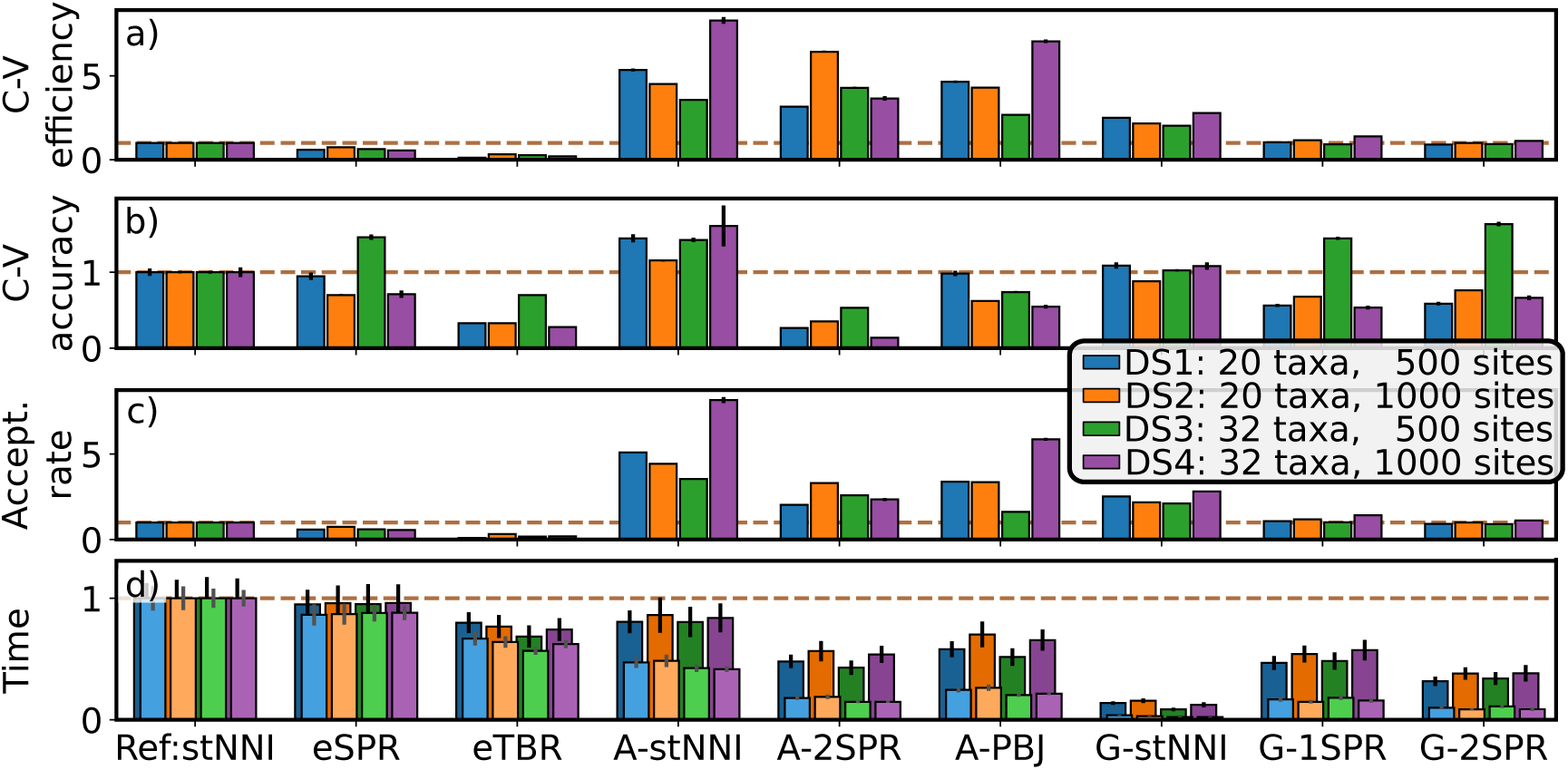
Performance improvements of proposals on simulated datasets using stNNI as reference. Figures a), b) and c) shows the relative increase in C-V efficiency, accuracy (1/distance) and acceptance rate, respectively. Figure d) represents the acceleration in computational time for the proposal (light colors) and for the whole MCMC iteration (proposal and likelihood evaluation; dark colors).

The performance of the naive proposals worsened proportionally to the amount of splits modified by a move, as shown by the decreases in all the metrics (Fig 7). Regardless of the proposal’s performance, using moves that alter multiple splits remains a mandatory feature for the estimation of tree distributions to ensure the proper exploration of the tree space. For instance, eSPR-type moves were able to visit tree topologies containing splits hardly reachable by stNNI-type proposals on dataset DS3; this advantageous feature was captured by the C-V accuracy (Fig. 7b for dataset DS3). The parsimony-guided proposals, as implemented in this study, performed poorly. The G-stNNI proposal was the only parsimony-guided proposal to improve the C-V efficiency by up to 2.5-fold factor, but at the cost of a 10-fold increase in computational cost.

The A-stNNI proposal revealed to be the best overall proposal by achieving consistent increases in C-V efficiency and C-V accuracy without leading to significant increases in computational cost. The adaptive N-edges proposals, the A-2SPR and A-PBJ presented different but complementary behaviors. The A-2SPR was very efficient at visiting a subset of the splits (specialist proposal), while the A-PBJ acted as a slightly less efficient N-edges version of the A-stNNI. These two adaptive proposals outperformed all their N-edges counterparts and therefore represent the best alternatives to the eSPR and eTBR proposals.

#### Mixtures performance and metric behaviors

The *mixed* and *best* mixtures converged toward the reference split frequencies as fast, if not faster, than the *naive* mixture (Fig. 8a). These mixtures, containing both type of proposals (guided and adaptive), were more consistent with respect to MCMC convergence than the mixtures composed of a single proposal type (e.g., *adaptive*). After removing the burnin, mixtures containing adaptive proposals accurately estimated the split frequencies 3- to 12- time faster than the *naive* and *guided* mixtures (Fig. 8b).

**Figure 8:**
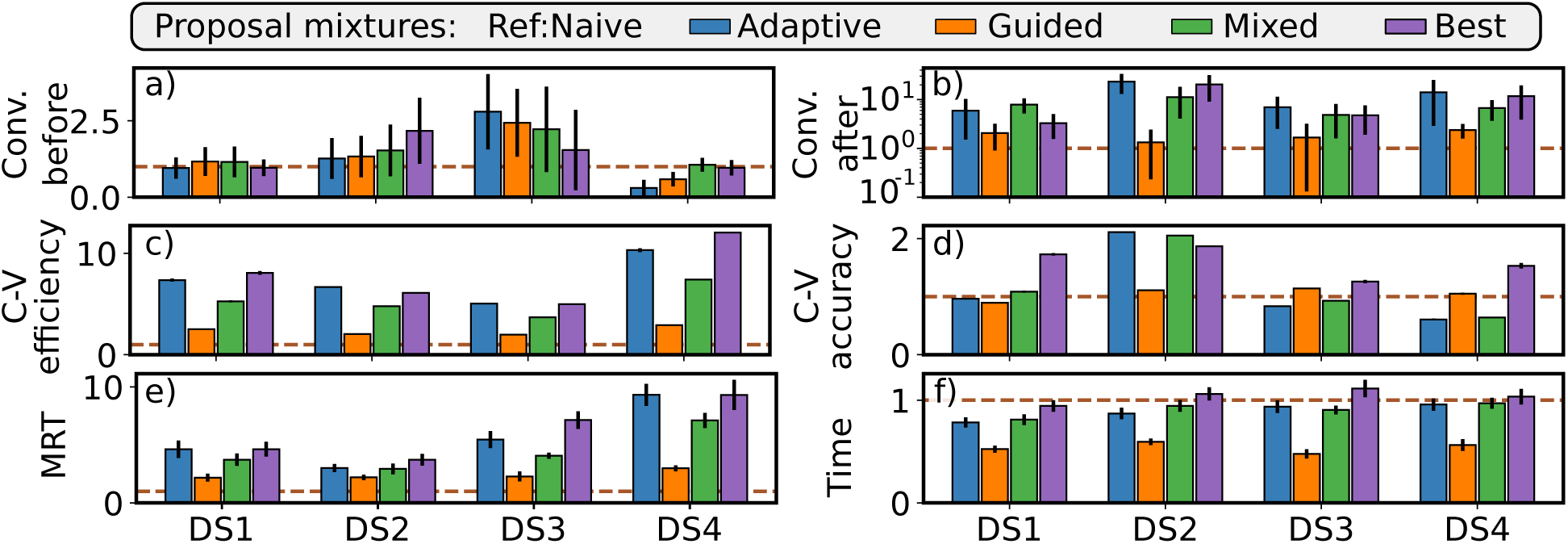
Performance improvements of proposal mixtures on simulated datasets using the naive mixture as reference. Figure a) and b) summarize the acceleration in convergence time before and after removal of the burnin phase, respectively. Convergence thresholds were fixed to a 2% and 1% average error per split. Figures c) and d) represent increases in C-V efficiency and accuracy (1/distance). Figures e) and f) illustrate the improvement in MRT and total runtime.

According to the C-V efficiency and the MRT metrics, the *adaptive* and *best* mixtures were up to 12 times more efficient at sampling the posterior distribution of trees (Fig. 8c and e). Their performance was closely followed by the *mixed* mixture but remained unequaled by the *guided* mixture. The C-V accuracy metric indicated that the *guided* mixture behaved similarly to the reference and that the *best* mixture was the only one to consistently perform as well or better than the *naive* mixture (Fig. 8d).

Most of the mixtures of proposals took slightly more computational time to reach 10 million iterations than the reference mixture. The *best* mixture was the only one to have a similar computational cost (Fig. 8f). The decreases in runtime observed on dataset DS2 and DS3 were due to a significant increase in acceptance rate of the adaptive proposals. The rejection of a proposal by the MCMC process induces computational procedures that restore the previous MCMC state. These procedures involve costly operations (i.e., memory backups or re-evaluations of likelihoods) that are avoided whenever a move is accepted. Lastly, the only mixture that significantly under-performed in term of runtime was the *guided* mixture due to the expensive computational cost of the G-stNNI proposal.

In addition to assessing the performance of the different proposal mixtures on simulated datasets, this experiment indicated that the C-V and MRT metric were significantly more stable than the convergence metrics. Convergence metrics are more likely to be impacted by the starting tree topology and stochastic behavior of the MCMC algorithm. More importantly, this experiment highlighted that the C-V efficiency and the MRT metrics were consistently measuring improvements having the same order of magnitude for all datasets and proposals.

### Proposal Mixture Performance on Empirical Datasets

The *best* proposal mixture consistently outperformed the *naive* mixture when considering the sampling of the posterior distribution of trees, regardless of whether MC^3^ was used (Fig. 9c and d). On challenging datasets, the use of the *best* mixture without MC^3^ increased the amount of successful MCMC runs but did not guarantee convergence within the imposed 10 million iterations (Fig. 9a). The *best* mixture with MC^3^ was the only setting under which convergence was consistently achieved. Overall, the *best* mixture without MC^3^ led to runs converging as fast as the *naive* mixture with MC^3^ and surpassed its performance when MC^3^ was not used (Fig. 9b). These improvements came at no significant additional computational cost: the variation in computational costs reached at most 11% of the reference cost, depending on the datasets and mixtures (SI Fig. 13).

**Figure 9:**
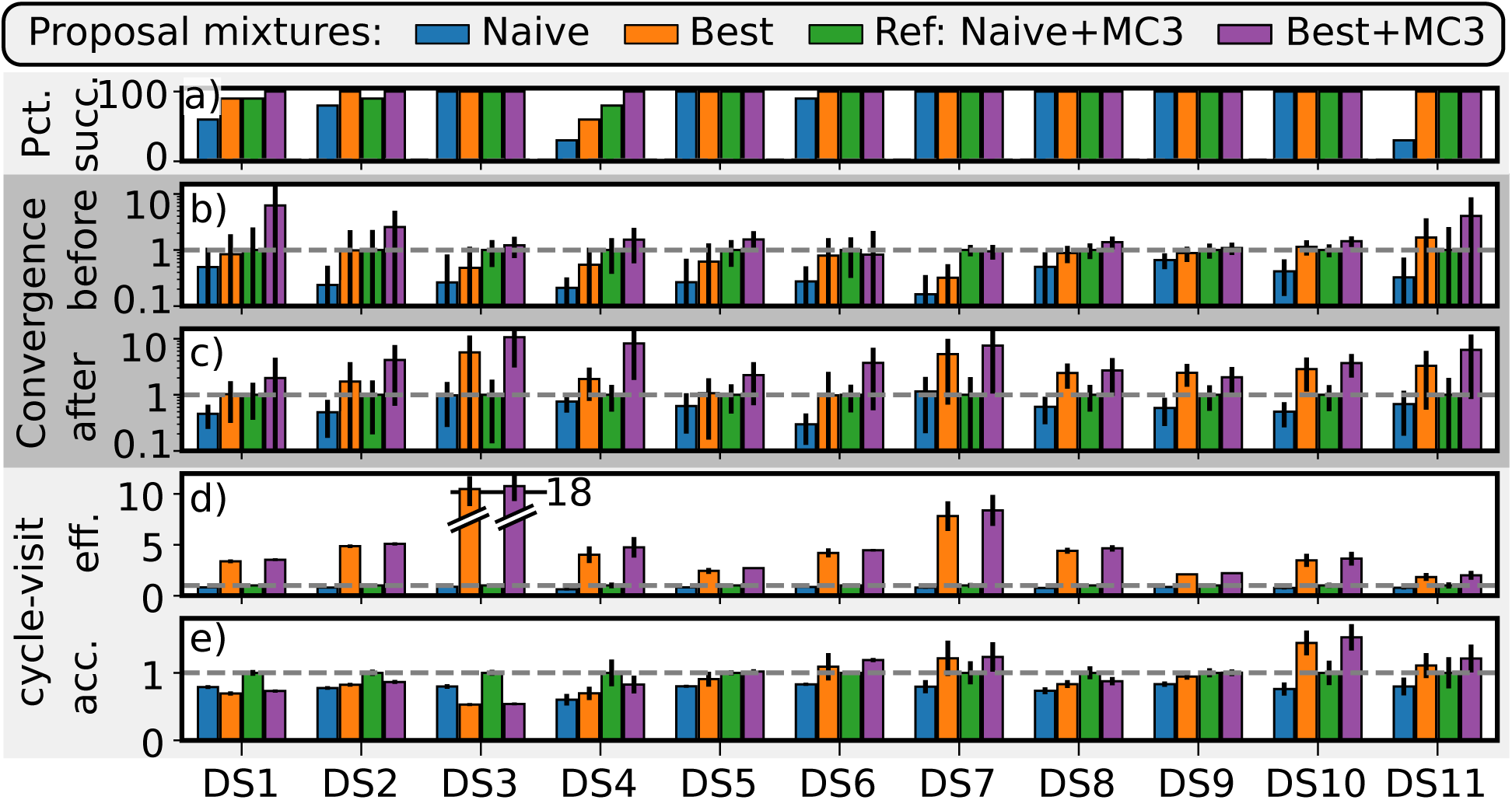
Performance improvements on empirical datasets using the *naive* mixture with MC3 as reference. Figure a) shows the percentage of runs having succeeded to converge within 10 million iterations. Figures b) and c) summarize the acceleration in convergence time before and after burnin removal, respectively. Convergence thresholds were fixed to a 2% and 1% average error per split. Figures d) and e) represent increases in C-V efficiency and accuracy.

**Figure 10:**
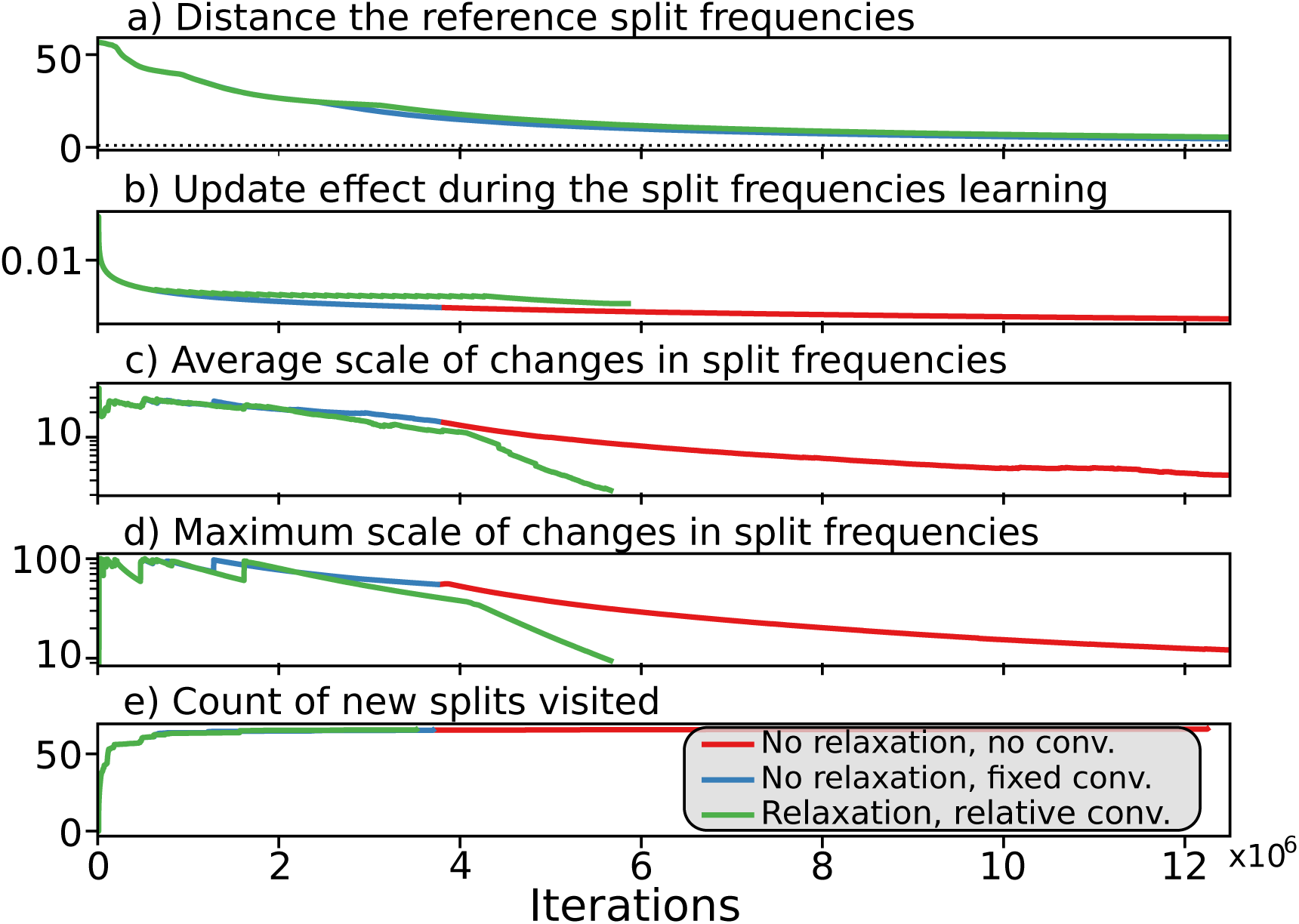
Learning process of the split frequencies.

Using the *best* mixture led on average to a 6-fold increase in C-V efficiency (with and without MC^3^) when compared to the reference mixture (*naive* plus MC^3^). The magnitude of the observed improvements were different depending on the datasets analyzed: the performance of the *best* mixture was directly correlated with the amount of information exploitable by the adaptive proposals in the distribution of split frequencies (SI, Fig. 11). Analyses on datasets with diffuse posterior tree distributions (i.e., DS5, DS9 and DS11) had only limited improvements in C-V efficiency that ranged from 2 to 2.7-fold.

**Figure 11:**
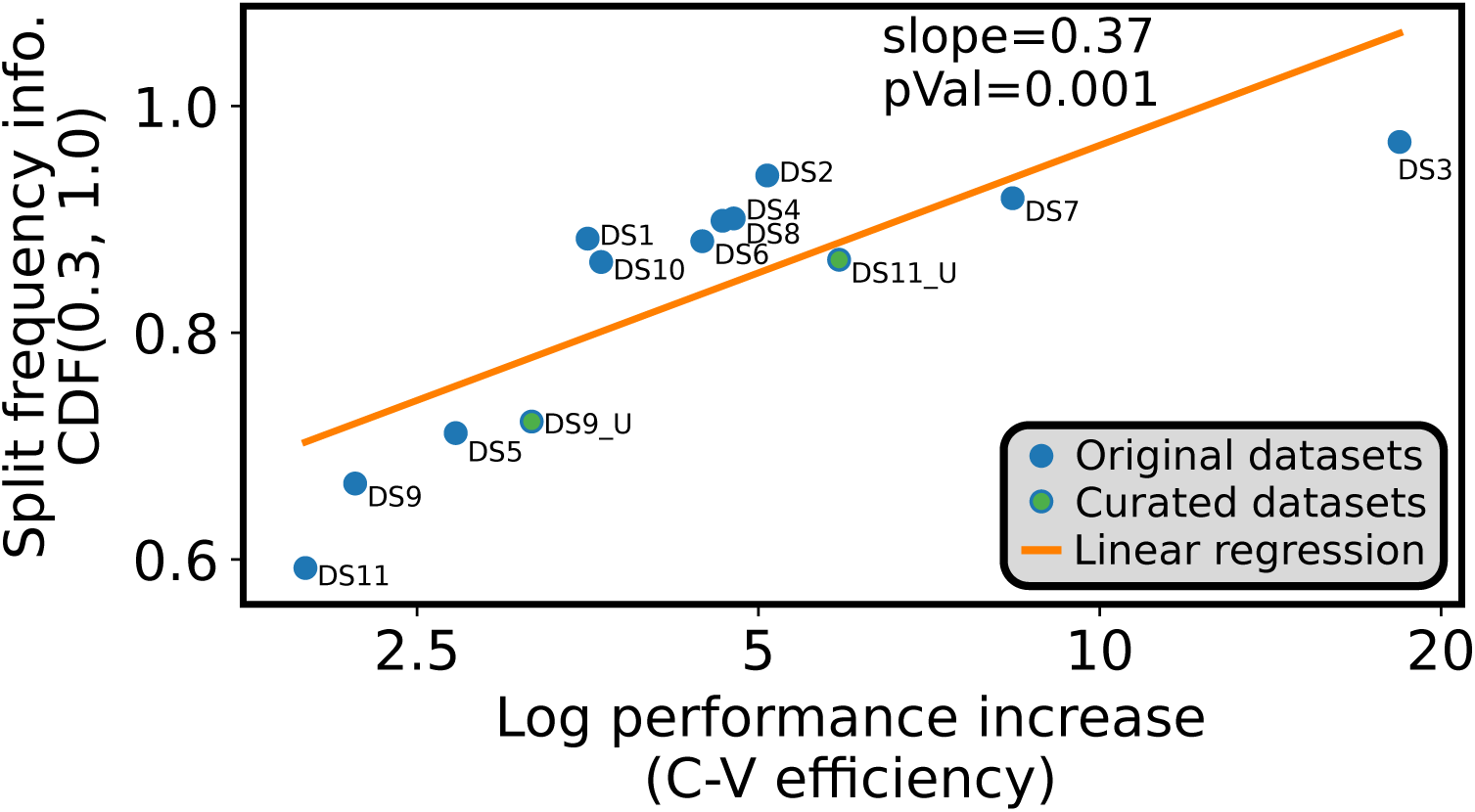
Linear regression of the information contained in the split frequencies and the log of the C-V efficiency increase. The information contained in split frequencies is computed as the CDF(0.3, 1.0) of the distribution of split frequencies and captures the amount of strongly structure in the tree.

Removing duplicated sequences from alignments DS9 and DS11 significantly increased their performance improvements (from 2.2 to 3.2 and 2 to 5.9-fold increases, respectively; SI, Fig. 11). For datasets with strong phylogenetic signal (e.g., DS3), the performance improvements reached up to an 18-fold efficiency increase. These observations could not be confirmed with the MRT metric due to its difficult application on diffuse tree topology distributions (Whidden and Matsen 2015). However, the observed trends of improvements and their magnitude concurred with the improvements in convergence speed after burnin removal (Fig. 9c).

Using the MC^3^ algorithm had a positive effect on the convergence of Markov chains regardless of the proposal mixture: all the runs converged with the *best* mixture and only a few failed with the *naive* mixture. These convergence failures happened on the three datasets (i.e., DS1, DS2, DS4) with the highest C-V distances (SI Fig. 12), indicating that many of their splits were difficult to visit. Nonetheless, when MC^3^ was used, convergence was reached on average 4.2 and 2 times faster before and after burnin removal, respectively. These improvements regarding the convergence were not captured by the C-V efficiency metric, indicating that the cold chain sampling efficiency remained mostly unaffected by MC^3^. The convergence improvements and small but systematic improvements in C-V accuracy resulting from the use of MC^3^ were however consistent with the effect of accepting bolder moves that could enable to explore different peaks of the posterior distribution (Fig. 9e).

**Figure 12:**
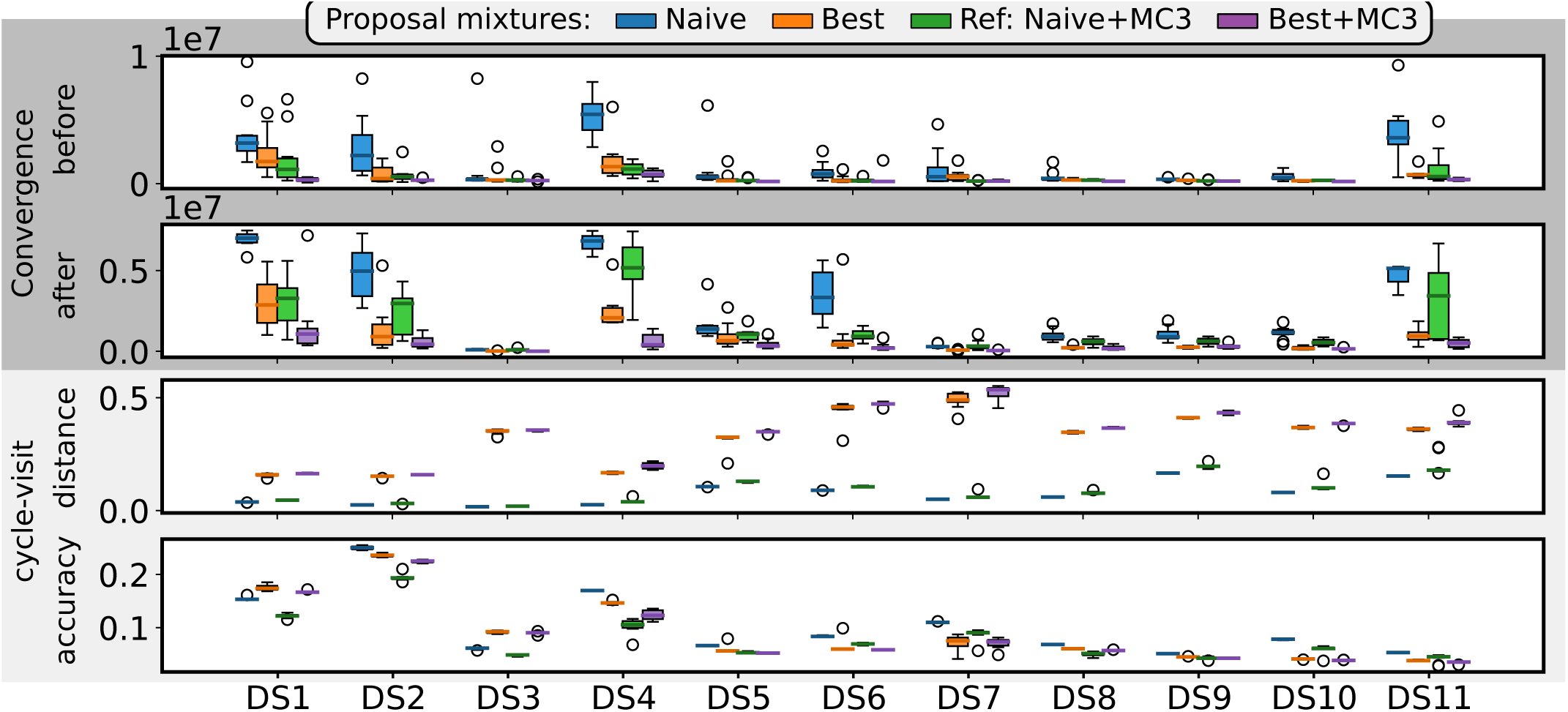
Absolute performance metrics on empirical datasets. Figure a) and b) summarize the convergence time before and after burnin removal, respectively. Convergence thresholds were fixed to a 2% and 1% average error per split. Figures c) and d) represents the C-V efficiency and distance.

**Figure 13:**
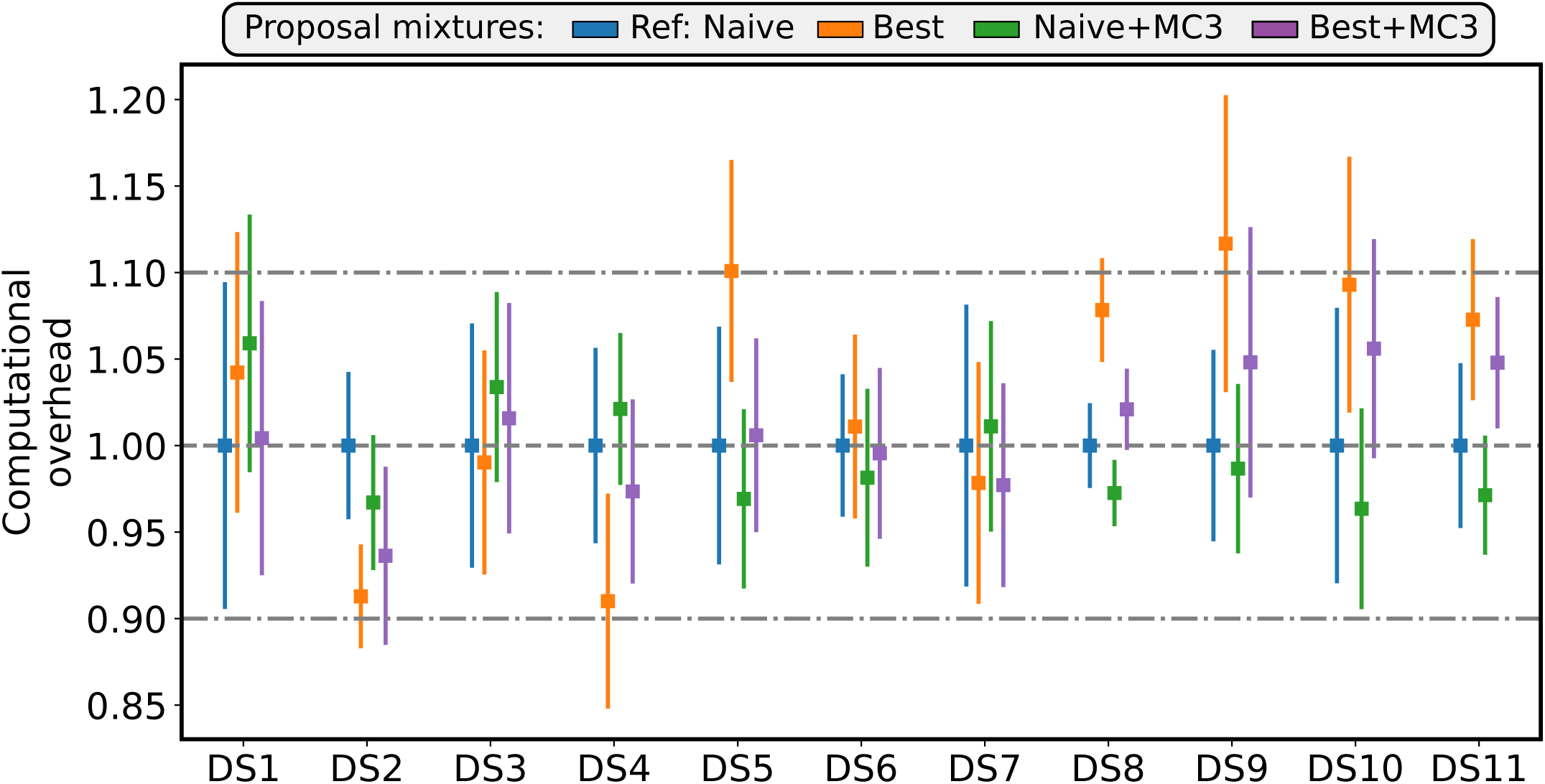
Overall computational overhead on empirical datasets using the naive mixture as reference. Analyses with MC3 were run using multiple processors (parallel).

When MC^3^ was not used, the C-V accuracy was improved by 22% on average by using the *best* mixture rather than the *naive* mixture.When MC^3^ was used, these performance improvements in C-V accuracy were however significantly different among the datasets and the proposals mixtures, and were not apparently correlated with the improvements in mixing efficiency (e.g., DS3). While adaptive proposals slightly improved the C-V accuracy, these observations suggest that their dominant effect was to increase the frequency at which appropriate moves were proposed, rather than enabling the visit of splits unreachable by naive proposal (e.g., multiple peaks).

## Discussion

In this study, I have presented the concept of adaptive proposals for unrooted tree topologies and developed a family of adaptive tree proposals. These adaptive tree proposals generate moves more likely to adequately sample the posterior distribution of trees by exploiting an estimate of the marginal split frequencies. This concept was applied to two standard proposals (i.e., stNNI and eSPR) generating specific type of moves. Additionally, I presented a novel philosophy for the design of tree proposals enabled by the concept of adaptive proposals.

This philosophy was used to design a proposal defining the most appropriate structure of moves by identifying strongly and weakly supported region of the phylogeny using a path-building mechanism. I showed that, while being more computationally expensive than standard proposal, the theoretic computational complexity of adaptive proposals was significantly lower than the complexity of parsimony-guided proposals and likelihood evaluations.

The performance of these adaptive tree proposals was assessed on simulated and empirical datasets. Using performance metrics designed for these experiments, I showed that adaptive proposals consistently outer performed their counterparts on simulated dataset. Using an empirically tuned proposal mixture to analyze 11 empirical datasets resulted in 2 to 18-fold improvements in mixing efficiency and up to 6 times faster convergence of MCMC and MC^3^ runs when compared to a standard proposal mixture composed of stNNI, eSPR and eTBR proposals. These performance improvements came without significant increase in computational cost and were correlated with the amount of phylogenetic signal in the alignments.

Adaptive proposals revealed to be superior to the naive and parsimony-guided proposals according to all metrics. The first key advantage enabling adaptive proposals to outperform other proposals is their ability to locate regions of a tree topology subject to uncertainties that could benefit from being modified. Non-adaptive proposals always start by arbitrarily choosing a region of the tree topology to modify. After this first arbitrary choice, naive proposals continue to apply randomized topological modification, while guided proposals use a score (e.g., parsimony or posterior probability Höhna and Drummond 2012)) to select the most promising resulting tree from a finite set of moves. Even if the guiding mechanism identifies good alterations, these proposals remain limited by the arbitrary choice made at first. I investigated an alternative approach for parsimony-guided proposals that involved an exhaustive exploration of all possible stNNI moves for a given tree. This proposal’s mixing efficiency was better than naive proposals but worse than adaptive proposals. Furthermore, its computational complexity exceeded the cost of a likelihood evaluation by an order of magnitude.

The second key advantage of adaptive proposals is their ability to exploit the *shape* of the posterior distribution of trees by cheaply approximating the split frequencies during an MCMC run. Contrary to guided proposals using proxy scores to approximate the posterior distribution (e.g., parsimony), adaptive proposals are not affected by the substitution models used for the analyses. However, this second key advantage is also the potential pitfall because the adaptive proposals strongly rely on accurate estimates of the split frequencies. Inaccurate estimates could lead adaptive proposals to have worse performance than their naive counterparts. When MC^3^ was used, this outcome was not observed on the 11 empirical datasets analyzed in this study, despite the ruggedness of their posterior distribution of trees (Whidden and Matsen 2015).

The concept of adaptive proposals as presented in this study is not limited to the three adaptive proposals that I developed, but opens new avenues toward the development of other tree proposals. For instance, the design philosophy used for the A-PBJ proposal that consists in designing proposals adaptively defining the move structure using the marginal split frequencies, could lead to other novel proposals by considering other path-building or structure-building strategies. Another avenue for improvements would be to estimate the joint distribution of split frequencies and exploit this information to consider bolder and more complex topological alterations. While the use of adaptive proposals could already benefit other types of phylogenetic inferences, these additional developments would be particularly beneficial to models having stronger constraints, such as clock-constrained or the inference of gene-trees within species trees (Rannala and Yang 2017).

In conclusion, the three adaptive tree topology proposals defined in this study represent a practical improvement for Bayesian inference of phylogeny, regardless of the substitution model considered. The concepts developed offer a fresh perspective on the design of tree proposals that should be advantageous to more challenging types of phylogenetic inferences and could therefore bring a new outlook to a challenging limitation existing since the early days of Bayesian inference of phylogenies.

## Acknowledgments

X.M. received funding from the Swiss National Science Foundation (P2GEP2 178032). I thank M. May and J. Huelsenbeck for discussion and feedback on this work.

## Supplementary Materials

### Learning Split Frequencies

To improve the learning of split frequencies, I used two frequencies per split: the internal and visible frequencies. The internal frequency is updated each *k*_1_ = 25 samples and is only internally used by the learning process. The visible frequency is the one used by the adaptive proposals and reflects the value of the internal frequency after *k*_2_ = 10 updates. Change of the split frequencies, as seen by the adaptive proposals, represents the behavior of the MCMC process averaged over a span of *k*_1_ × *k*_2_ = 250 iterations.

To improve the rate at which split frequencies can vary, I uses a relaxation mechanism that can be triggered two ways: upon an increase of 15% in the total number of splits encountered during the run or upon a significant change in the estimated marginal split frequencies. Changes in split frequencies are assessed by monitoring the trends of change of each split frequency over the last *k*_3_ = 1000 updates. The trend of change is estimated as the ratio between the absolute value of the observed change over *k*_3_ iterations and the maximum expected change over the same span of iterations. A relaxation is triggered whenever the average observed trend of change (averaged over all splits) exceeds 10% or if the maximum observed trend of change exceeds 30%. A trigger of the relaxation mechanism enables the estimates of the split frequency to adapt faster by multiplying the splits counts *cnt*(*e*_*i*_) and the total count *cnt* by a factor *max*(0.9, *k*_4_/*cnt*). To ensure stability, a relaxation can only be triggered once every *k*_5_ = 20 × *k*_1_ updates and can not reduce *cnt* under *k*_6_ = 2000.

After a significant period of stability in split frequencies, the learning process is terminated and the frequencies are fixed. The convergence is detected once *cnt* reach *k*_7_ = 10000. The convergence is therefore achieved once there has been no relaxation for a significant amount of iterations, implying that the number of new splits is not subject to significant increase and that the scale of change in split frequencies is low.

Figure 10 illustrates the effect of the relaxation mechanism and the relative convergence detection on a simulated dataset. The relaxation mechanism when active maintains the scale of change per update until the trends of change per split frequency reach their thresholds (Fig. 10b, c and d). The relative convergence termination enables then the split frequencies to continue their convergence for several iterations before halting the learning process. Runs without the relaxation mechanism reveal two problematic behaviors: the run with fixed convergence threshold terminate while split frequencies are still being learned, while the run without convergence threshold shows that the split frequencies are still being consistently altered after 1 million iterations (Fig. 10).

### Stochastic Component

The estimation of the split frequency may turn out to be unreliable, specifically during the early steps of the learning process. I therefore augment the adaptive tree proposals with a stochastic component based on a residual term *ϵ* drawn from an exponential distribution with parameter *λ*. The value of *λ* follows a decreasing schedule (e.g., from 5.0 to 0.01) during the earliest steps of the learning process. This strategy improves adaptive proposals in two ways. First, moves proposed when the split frequencies are unreliable are acting similarly to naive moves, while still accounting weakly for split frequencies. As these estimates stabilize, the effect of the stochastic component decrease but continues to enable the consideration of moves unsupported by the split frequencies. Proposing unsupported moves remains advantageous by enabling the MCMC to visit unexplored parts of the tree space or by limiting the effect of inaccurate split frequency estimates.

### Parsimony Score Transformation

While the concept of using different scores to guide a tree proposal is well defined (Höhna and Drummond 2012), there exists to my knowledge no reference on how to convert a parsimony score into a valid probability. The transformation procedure considers proposals that, given tree *τ*, enumerates *K* moves leading to trees 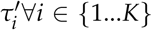 and uses the function *h*(·) to compute the parsimony score of a tree (e.g. fast-Fitch algorithms (Ronquist 1998)).

I first re-scale the relative difference between the parsimony score of the current tree *h*(*τ*) and the one of each of the proposed trees 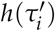 to fit within the [−1..1] interval according to

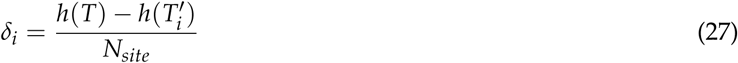

The extreme values of the [−1..1] interval are unrealistic to be encountered for *δ*_*i*_ on local moves affecting at most less than a tenth of splits. The *δ*_*i*_ scores are therefore truncated within the [−0.1. 0.1] interval and linearly transformed to fit within the [0..1] interval according to

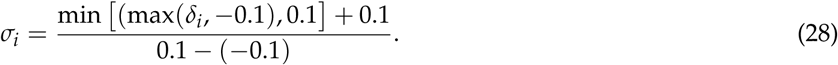

The probability of proposing move 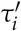 is then defined as

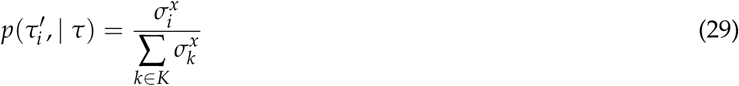

where *x* can be used to tune the behavior of the move ranging from a naive behavior with *x* = 0, to a strongly guided proposal with *x* ≫ 0. I set *x* to 8 based on empirical observations.

### Computational Complexity of Proposals

The theoretical computational complexity of the different type of proposals provides a good indication of their computational overhead with respect to the evaluation of a likelihood. An alignment contains the molecular state of *m* sites for *n* taxa, where the state is defined over *k* characters (e.g., *k* = 4 for nucleotide sequences). The theoretical cost of computing a parsimony score is of *O*(*mnk*) operations, while the one of a likelihood using the Felsenstein pruning algorithm costs *O*(*mnk*^2^). To account for the use of partial evaluations (i.e., refreshing computations on a subset of the tree), I use the 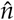 notation defining a subset of operations on *n*. Similarly, the use of the Gamma model for rate heterogeneity is accounted for using the *c* variable to define the number of rate categories, leading to a complexity of 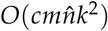 for a partial likelihood evaluation.

A naive tree proposal (e.g. stNNI or eSPR) has a computational complexity proportional to the number, *s*, of splits it alters (e.g. *O*(1) for the stNNI or *O*(*s*) for the eSPR). Parsimony-guided proposals have a complexity proportional to the size of the set of moves generated and the cost of estimating parsimony score for each of them. The *G* −*NSPR* generates *O*(2^*s*^) moves, where *s* represents the maximum number of edges, for the eSPR move, while the *G* − *stNNI* estimates 2 moves for each internal edge (*O*(*n*)) resulting in proposal complexity of 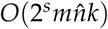 and 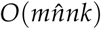, respectively.

The complexity of adaptive tree proposals is generally proportional to the size of the tree (≈*O*(*n*)). More specifically, the A-stNNI proposal requires the computation of the normalization constant for the internal edge probabilities (denominator of Eq.(3)) that costs *O*(*n*). The A-2SPR proposal requires the enumeration of all *s*-edges contiguous paths with *s* = 2 and the normalization of their probabilities (*O*(2^*s*^*n*)). The enumeration of the 8 possible moves and the estimation of the change in split frequencies over *s*-edges costs *O*(8*s*), leading to a total complexity of *O*(*s* + 2^*s*^*n*). The A-PBJ proposal requires the calculation of all internal edge probabilities (*O*(*n*)) and the construction of three paths. Unstable paths cost each ≈*O*(*s*_1_) and ≈*O*(*s*_2_), respectively. The stable paths 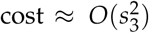 due to the marginalization over all possible central edges (Eq.(19)). This proposal’s total complexity is of the order of 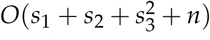. While each path has a different length *s*, the sum of the path sizes has for constraint that *s*_1_ + *s*_2_ + *s*_3_ *< n* given that the concatenation of all three paths represents a contiguous path in the tree. The complexity of the A-PBJ, under realistic settings, is therefore at most 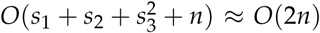. Extreme cases could lead this complexity to ≈ *O*(*n* + *n*^2^), but would imply a stable path including nearly all internal edges of the tree. Only nearly linear bifurcating trees having each posterior probability of split frequencies close to 1 would induce this type of therefore improbable scenario.

### Dataset Simulation Setting

I simulated dataset with 20 and 32 taxa. For each taxa number, I simulated alignment composed of 500 and 1000 sites. The tree topology and branch length of these 4 datasets were simulated under a birth death model with 1.9 speciation rate, 1.1 extinction rate and the tree length was rescaled such as to have an expected branch length of 0.1. Alignments were simulated under a GTR model with realistic stationary frequencies (*π*_*ACTG*_ = (0.3, 0.2, 0.2, 0.3) and exchangeability rates (*a* = 0.06, *b* = 0.11, *c* = 0.28, *d* = 0.06, *e* = 0.33, *f* = 0.17) that were estimated from a 16S ribosomal RNA alignment.

### Prior Settings

Simulated datasets were analyzed under a GTR model, while empirical datasets were analyzed under a GTR+Gamma model with 4 discrete rate categories. I used flat Dirichlet priors for the state frequencies and the exchangeability rates. Exponential distributions with rates *λ* = 10 were assigned to branch lengths and with rate *λ* = 1 to the *α* parameter of the Gamma model.

